# Forty New Genomes Shed Light on Sexual Reproduction and the Origin of Tetraploidy in Microsporidia

**DOI:** 10.1101/2025.05.12.652816

**Authors:** Amjad Khalaf, Chenxi Zhou, Claudia C Weber, Emmelien Vancaester, Ying Sims, Alex Makunin, Thomas C Mathers, Dominic E Absolon, Jonathan MD Wood, Shane A McCarthy, Kamil S Jaron, Mark Blaxter, Mara KN Lawniczak

## Abstract

Microsporidia are single-celled, obligately intracellular parasites with growing public health, agricultural, and economic importance. Despite this, Microsporidia remain relatively enigmatic, with many aspects of their biology and evolution unexplored. Key questions include whether Microsporidia undergo sexual reproduction, and the nature of the relationship between tetraploid and diploid lineages. While few high-quality microsporidian genomes currently exist to help answer such questions, large-scale biodiversity genomics initiatives, such as the Darwin Tree of Life project, can generate high-quality genome assemblies for microsporidian parasites when sequencing infected host species. Here, we present 40 new microsporidian genome assemblies from infected arthropod hosts that were sequenced to create reference genomes. Out of the 40, 32 are complete genomes, eight of which are chromosome-level, and eight are partial microsporidian genomes. We characterised 14 of these as polyploid and five as diploid. We found that tetraploid genome haplotypes are consistent with autopolyploidy, in that they coalesce more recently than species, and that they likely recombine. Within some genomes, we found large-scale rearrangements between the homeologous genomes. We also observed a high rate of rearrangement between genomes from different microsporidian groups, and a striking tolerance for segmental duplications. Analysis of chromatin conformation capture (Hi-C) data indicated that tetraploid genomes are likely organised into two diploid compartments, similar to dikaryotic cells in fungi, with evidence of recombination within and between compartments. Together, our results provide evidence for the existence of a sexual cycle in Microsporidia, and suggest a model for the microsporidian lifecycle that mirrors fungal reproduction.

## Introduction

Biology is characterised by intimate interactions and a fundamental interdependence between organisms living in close proximity. Yet, our understanding of symbionts and cobionts (organisms sampled alongside a target organism) typically lags behind our understanding of their hosts. Primarily, this is due to difficulties in accessing an organism’s symbionts, growing them, and/or assessing their behaviour. One such example is Microsporidia, which are single-celled, spore-forming, obligately intracellular parasites [1]. They were first described as the agent of a disease known as “pébrine” in farmed silkworms (*Bombyx mori*, Lepidoptera), which had caused crises in the industry along the silk road in the late 1800s [2,3]. Since then, Microsporidia have been identified in many different hosts, and are now appreciated as parasites of a broad range of metazoans, and even some protozoans [4,5]. Nine genera have been identified as human pathogens, especially in immunocompromised individuals, causing a range of symptoms such as diarrhea, encephalitis, keratitis, sinusitis, and myositis [6–10]. Similarly, arthropod-infecting microsporidians constitute a growing problem for beekeeping and aquaculture industries worldwide [11–14]. Other microsporidians are being explored as potential malaria transmission control agents, with evidence associating infection of *Anopheles* mosquitoes to a reduction in *Plasmodium* transmission [15–17].

Despite their importance, microsporidian biology remains fairly enigmatic, owing in part to their obligately intracellular nature, small size, low prevalence in most host populations, and biological quirks such as cryptomitosis (where the condensation and separation of chromatin into distinct chromosomes are unclear) [4,18,19]. One oddity that some microsporidian species share with diplomonads such as *Giardia intestinalis* is the persistence of two equivalent nuclei inside one cell, closely appressed against each other with distinct nuclear membranes and synchronous replication, for the whole lifecycle, a form known as a “diplokaryon” [20–23]. Some microsporidians remain monokaryotic for all their lifecycle [24], and others cycle between diplokaryotic and monokaryotic forms [21,25–28].

The cycling between monokaryotic and diplokaryotic states in some microsporidian species has widely been assumed to be part of a meiotic reproductive cycle, and phenomena interpreted as gametogenesis, plasmogamy, karyogamy, and synaptonemal complexes have been reported [21,24,29–40]. Whilst strictly diplokaryotic and strictly monokaryotic species were assumed to only undergo asexual mitosis [41], recent observations such as a very transient monokaryotic stage and synaptonemal complexes in diplokaryotic species suggest otherwise [42,43]. Population-level genetic data has indirectly suggested the occurrence of rare or recent recombination events in two strictly monokaryotic species [44,45]. Although the presence of meiosis has never been validated beyond morphological data, homologues of genes involved in meiosis have been identified in many microsporidian species, with phylogenetic analysis indicating that all Microsporidia may have descended from a sexual fungal ancestor [18]. Furthermore, the ploidy of each microsporidian nucleus, and whether that changes during developmental or meiotic cycles, remains an open question. Seven microsporidian species have been reported to be tetraploid, including both monokaryotic and diplokaryotic species [46–48]. Additionally, some microsporidians have species-specific lifecycle variants, such as the formation of an octet of spores as one infective unit (an “octospore”) [29,49], and it is unknown whether the ploidy of these is the same as their monokaryotic/diplokaryotic forms.

Ploidies other than haploid and diploid are common in nature but generally phylogenetically unstable, and are usually resolved back to effective diploidy through loss of one set of chromosomes or genome-wide rediploidisation leaving the signatures of whole genome duplication [50]. The high frequency of apparent tetraploids in Microsporidia [46] could be a reflection of a tendency towards formation of tetraploids or a reflection of a fundamental tetraploid state. It is unclear whether observed microsporidian polyploids arose through autopolyploidy, where the four genome copies derive from within-species events, or allopolyploidy, where genomes from different species are combined through hybridization (reviewed in [51]). Furthermore, it is unknown whether tetraploidy in Microsporidia is characterised by a single ancient event that has been stably maintained across multiple lineages, an outcome that would be highly unusual given that rediploidisation is typically a key process that follows whole-genome duplications [52–60], or whether polyploidy has arisen independently in each microsporidian lineage where it is observed.

While the number of microsporidian genome assemblies is increasing, few are high quality, chromosome-level assemblies. High quality assemblies are crucial to disentangling questions about the occurrence of sexual reproduction, the origins of polyploidy, and the interaction of the two [61]. Large-scale reference genome and next-generation sequencing initiatives, such as the Darwin Tree of Life (DToL) [62], can incidentally generate high-quality genome assemblies for cobionts when sequencing host species, and thus offer an unrivaled data-generation opportunity for rare and unculturable endosymbionts. For instance, DToL has recently released over 100 novel *Wolbachia* genomes and two cnidarian endoparasite genomes assembled from data arising from individual hosts targeted for reference genomes [63,64]. Some screens for Microsporidia have also been carried out on DToL data, yielding a few microsporidian genome sequences [61,65,66].

Here we present microsporidian genomic data from 40 host organisms sequenced at the Wellcome Sanger Institute as part of DToL. We recover eight partial and 32 complete (or nearly complete, with Benchmarking Using Single Copy Orthologues [BUSCO] completeness scores >70%) microsporidian genomes. Eight of our complete genomes are chromosome-level assemblies, seven of which, to our knowledge, were scaffolded with the first Hi-C data generated for Microsporidia. These new genomes represent much of the breadth of currently described microsporidian diversity, with genomes from five of the seven microsporidian clades named by Bojko and colleagues [4]. We show that one tetraploid genome is organised into two compartments, likely the nuclei of the diplokaryon, but also show evidence of historical recombination both between all four genomes. We describe rearrangements between the haplotypes of some genomes, chromosomal rearrangements in microsporidian evolution, and a high tolerance for segmental duplications. We recognise recombination signatures in other tetraploid genomes, and propose a model to synthesise our observations of ploidy, reproduction and the microsporidian lifecycle.

## Results

### Forty new microsporidian genome assemblies

We identified host organisms carrying microsporidian infections by screening the raw genome sequencing data generated by DToL for microsporidian sequences, or by PCR amplification of microsporidian targets from host DNA extracts before genome sequencing. We also observed microsporidian-like sequences in additional genomes that proved to be horizontal DNA transfers into the host genome in the absence of current live infections. From 40 host species, we recovered 32 complete microsporidian genome assemblies with BUSCO completeness score >70%, including eight chromosome-level genomes, seven of which were scaffolded with Hi-C data. We also assembled eight partial genome sequences, with BUSCO completeness score <70%. The recovered microsporidian genomes come from hosts belonging to eight insect orders, with lepidopteran and dipteran hosts yielding 13 and 12 microsporidian genomes, respectively (Fig. 1). The hosts include several not previously known to be infected by microsporidia, including *Loensia variegata* (Psocodea, bark lice), *Vulgichneumon bimaculatus* (Hymenoptera, ichneumon wasp), and *Delia platura* (Diptera, seedcorn maggot fly) (Supporting Information Section 1). Our sample size is small, but we note that most microsporidian genomes from lepidopteran hosts are derived from the microsporidian group Nosematida (77%, n = 10), and most microsporidian genomes from dipteran hosts are derived from Amblyosporida (67%, n = 8). While some microsporidians are known to distort the sex of their host populations [67], the sex of our genomes’ hosts was unknown in most cases (24 species). In the remaining cases, nine were identified as female and seven as male (Fig. 1). A relatively equal proportion of female and male hosts are infected with Nosematida (Fig. 1), but we could not assess skews in host sex ratios for other microsporidian groups due to missing data on sex (for Amblyosporida-infected hosts), or a small sample size (for Neopereziida-infected hosts) (Fig. 1). The full list of hosts and their recovered microsporidian genome assemblies is given in Supporting Information Section 1.

**Fig. 1:**
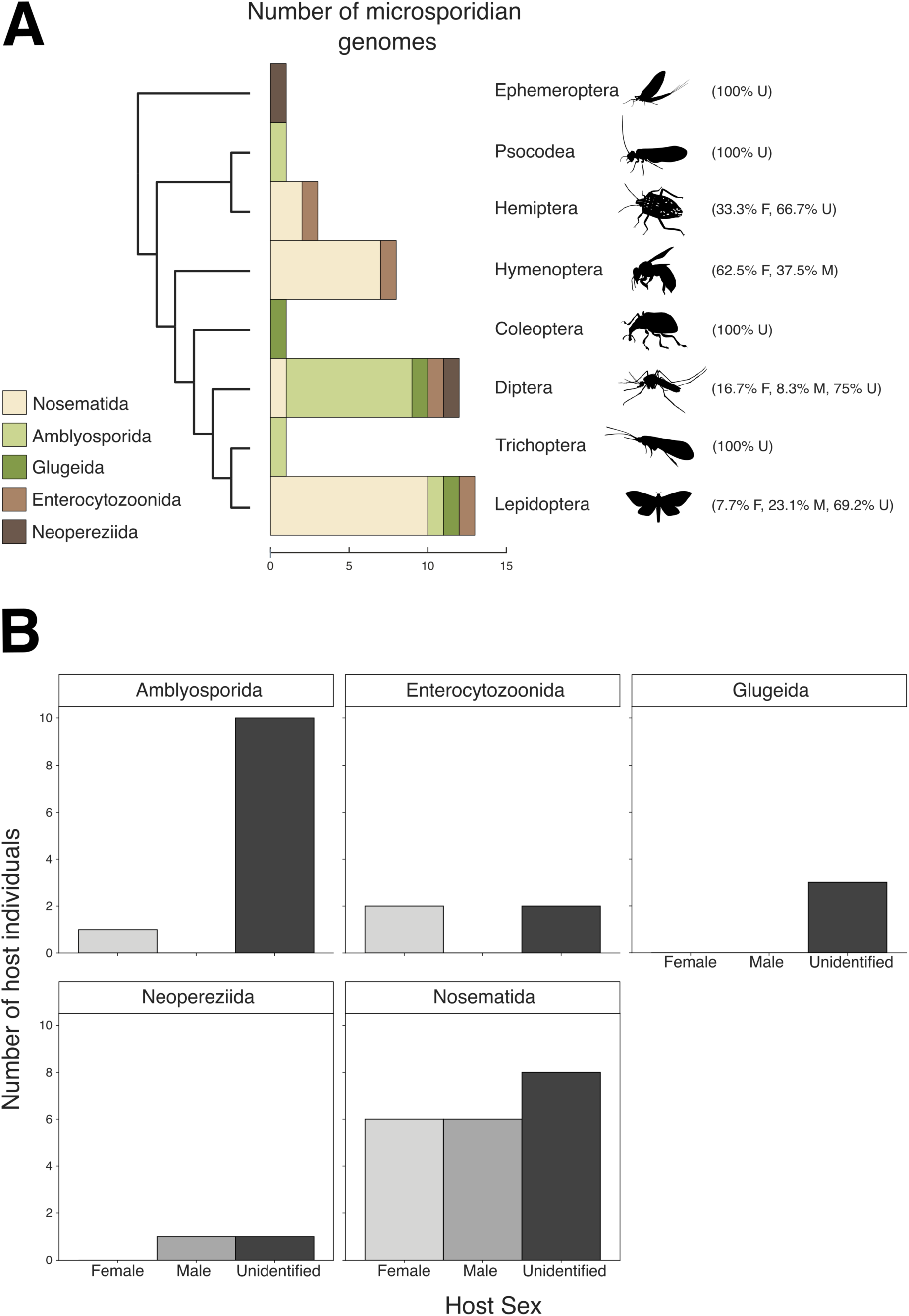
Prevalence of Microsporidia in DToL insect genomes. Microsporidian genomes recovered from insect hosts, split by taxonomic order in (A) and by host sex in (B). F: female, M: male, U: unspecified sex. Credits for the silhouettes used in this figure (https://www.phylopic.org): Ephemeroptera, Nathan Jay Baker; Psocodea, Graham Montgomery; Hemiptera, Dave Angelini; Hymenoptera, Richard Harris; Coleoptera, Kanako Bessho-Uehara; Diptera, T. Michael Keesey; Trichoptera, Hester Weaving; and Lepidoptera, T. Michael Keesey.

The genome sequences were placed in five of the seven microsporidian clades identified by Bojko and colleagues [4]. Half derived from Nosematidae (two chromosome-level genomes, 13 complete genomes, five partial genomes) and 11 from Amblyosporidae (six chromosome-level genomes, five complete genomes) (Fig. 1 and Fig. 2). Four belonged to Enterocytozoonidae (one chromosome-level genome, two complete genomes, one partial genome), two to Neopereziidae (two partial genomes), and three to Glugeidae (three complete genomes) (Fig. 1 and Fig. 2). Most chromosome-level and complete genome assemblies presented here have comparable or higher contiguity and BUSCO completeness compared to previously published microsporidian genome assemblies (Fig. 2). Our complete genome sequences range in span from 2.35 Mb to 56 Mb. The eight largest genome assemblies are unpurged and thus likely retain some haplotypic duplication, while the largest purged assembly (idChiSpeb1.µ, see Materials and Methods for an explanation of the naming system used) is 20 Mb (Fig. 2). This falls within the range of previously sequenced microsporidian genomes, which range in size from 2.2 Mb (*Encephalitozoon romalae*) to 51.3 Mb (*Edhazardia aedis*) [68].

**Fig. 2:**
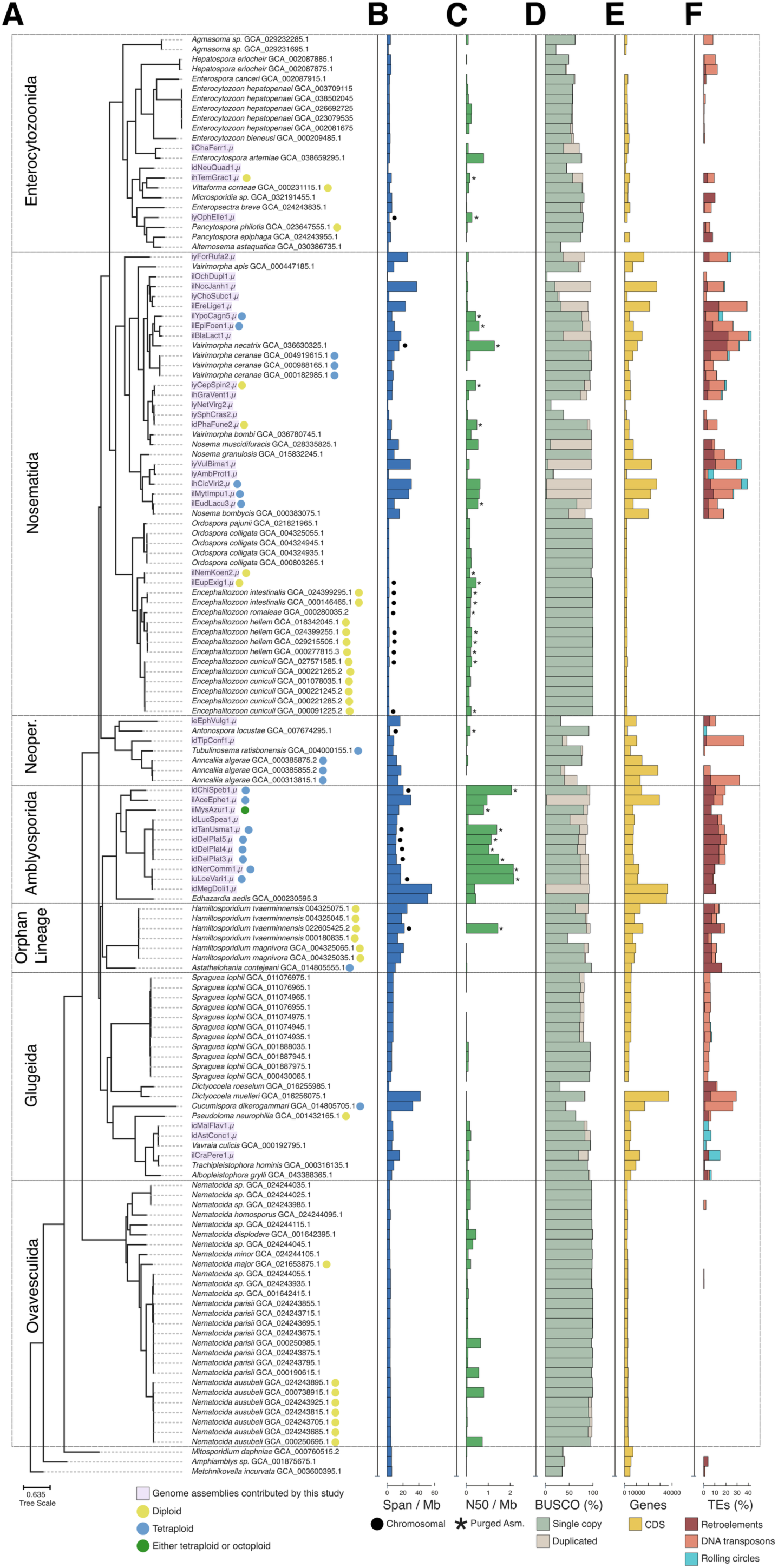
600 Gene Phylogeny of Microsporidia. (A) ASTRAL [69] phylogeny summarising individual phylogenies of 600 BUSCO genes (microsporidia_odb10) [70] across all publicly available microsporidian genome assemblies (n = 106), and the genome assemblies generated in this study (n = 40, marked in purple). Branch lengths were estimated with IQ-TREE using a concatenated alignment of the individual BUSCOs [71]. Ploidy is marked in circles at the tips of the tree for genomes where it was characterisable. (B) Genome assembly span (Mb), with black circles marking chromosome-level genome assemblies. (C) N50 values (Mb), with asterisks marking purged genome assemblies. (D) BUSCO gene (microsporidia_odb10) completeness percentage, marked in green for single-copy genes, and beige for duplicated genes. (E) Gene count as predicted by Prokka [72]. (F) Transposable element percentage as predicted by RepeatModeler and RepeatMasker [73,74], marked in burgundy for retroelements, peach for DNA transposons, and blue for rolling circles.

We performed gene annotation (with Prokka) and repeat annotation (with RepeatModeler and RepeatMasker) on all genomes [72–74] so that all genomes were annotated congruently. To assess the relationship between the phylogeny and the number of coding sequences, transposable element loads, and genome spans, we computed transformations representing the fit of each feature with the tree’s topology (λ), branch-lengths (κ) and root-tip distance (δ) [75] (Supporting Information Section 2). For the majority of examined traits, we found no significant phylogenetic signal, with the exception of a strong root-tip distance effect on all transposable element loads. Genome span was strongly correlated with the number of coding sequences (Supporting Information Section 2). Retroelement load and DNA transposon loads were moderately correlated to one another and genome span, whereas helitron load was weakly correlated to all other features.

The phylogenetic tree suggests a revision of microsporidian relationships. We confidently placed Nosematida as a sister group to Enterocytozoonida, and the group known as ‘orphan lineage’ (containing the genera *Hamiltosporidium* and *Astathelohania*) as a sister group to Amblyosporida, in agreement with previous whole-genome phylogenies [4,76,77]. However, Glugeida was a sister group to the ancestor of the orphan lineage and Amblyosporida. The previously contested place of Neopereziida as a sister group to the ancestor of Nosematida and Enterocytozoonida was confirmed (Fig. 2).

The classification of species traditionally assigned to *Nosema*, such as *Nosema* (=*Vairimorpha*) *ceranae* and *Nosema* (=*Vairimorpha*) *apis*, has been the subject of ongoing debate [4,47,78,79]. Our phylogeny, using 600 loci, strongly supports the split between *Vairimorpha* and *Nosema*, with *Nosema* (=*Vairimorpha*) *ceranae* clustering robustly with other *Vairimorpha* genomes (Fig. 2).

### Operational taxonomic unit classification of species suggests autotetraploidy in Microsporidia

Morphological and histopathological data are usually employed to identify microsporidia to species level, but no such data were available for these newly assembled microsporidian genomes. We measured phylogenetic branch lengths between every possible combination of two genomes to establish a baseline for genomic disparity among individuals that are known to be members of the same species based on morphological, histopathological, or cell culture data. We then used this baseline to assess whether any of the new genomes could be diagnosed as members of known species, and whether the homeologous subgenomes of our tetraploid genomes could be characterised as belonging to the same species (autotetraploidy) or not (allotetraploidy).

Publicly available genomes classified to the same species exhibited high genomic similarity, with an average branch length distance of 0.01 amino acid substitutions per site (ranging from 0 to 0.032 amino acid substitutions per site, estimated from BUSCO genes). For example, *Encephalitozoon hellem* genomes were separated by branch lengths of 0 to 0.015 amino acid substitutions per site, *Nosema* (=*Vairimorpha*) *ceranae* genomes by 0 to 0.003 amino acid substitutions per site, and *Nematocida ausubeli* genomes by up to 0.023 amino acid substitutions per site (Fig. 3). Genomes from different species were much more divergent, with an average branch length of 0.577 amino acid substitutions per site. One pair of species, *Hamiltosporidium tvaerminnensis* and *Hamiltosporidium magnivora*, had particularly short branch lengths (0.012–0.017 amino acid substitutions per site). Excluding these, the smallest branch length between different species within the same genus was 0.15 amino acid substitutions per site.

**Fig. 3:**
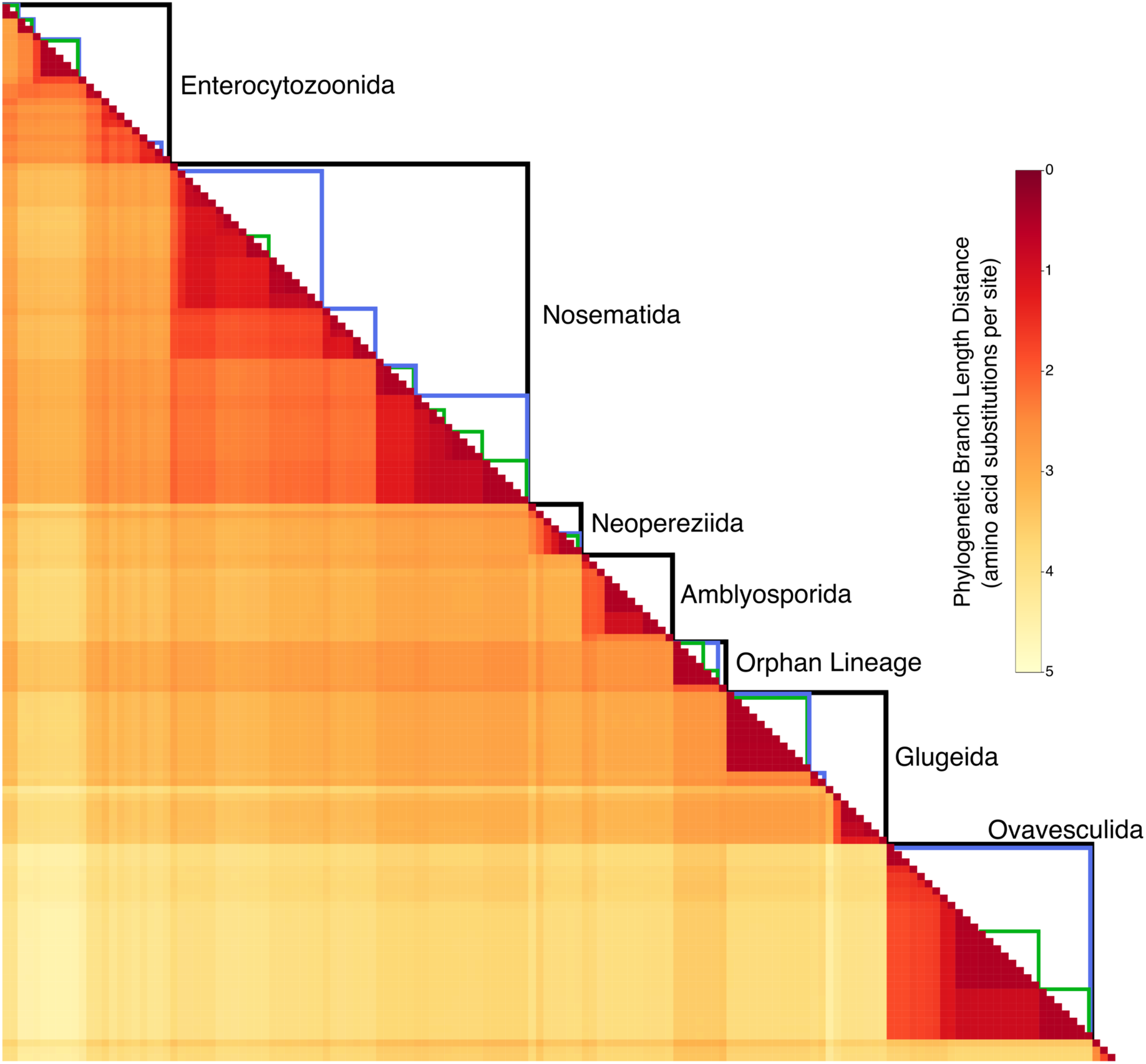
Pairwise phylogenetic branch lengths between genomes. The heatmap shows the phylogenetic branch length distance (in amino acid substitutions per site) for every combination of two genomes from the complete genomes generated in this study and all publicly available genomes, ordered by the phylogeny in Fig. 2. Black lines delimit clades named by Bojko and colleagues [4], and blue lines delimit genera. Green lines delimit genomes characterised as belonging to the same species using morphological or histopathological data.

Relying on these data, we established a conservative branch length threshold to classify genomes as belonging to the same species based on the smallest distance between *H*. *tvaerminnensis* and *H*. *magnivora* genomes (0.012 amino acid substitutions per site). Using this criterion, we classified 17 of the new genomes as belonging to a known species or an unnamed species formed by two or more of our genomes (Supporting Information Section 3). If within-species divergence was permitted to extend to the full range observed within species (i.e. to 0.032 substitutions per site), one additional new genome was classified as being a likely member of a named species. From these analyses we identified five species-like groupings of otherwise unidentified microsporidia sequenced here (Supporting Information Section 3). Those include gmOTU1 comprising iuLoeVari1.µ (from host *Loensia variegata* [Psocodea])), idNerComm1.µ (from host *Neria commutata* [Diptera)] and idMegDoli1.µ (from host *Megamerina dolium* [Diptera]); and gmOTU2 from five dipteran hosts: idDelPlat3.µ (from host *Delia platura* [Diptera]), idTanUsma1.µ (from host *Tanytarsus usmaensis*), idDelPlat4.µ (from host *Delia platura*), idLucSpea1.µ (from host Lucilla sp.) and idDelPlat5.µ (from host *Delia platura*). For an explanation of the naming system used for OTUs, see Materials and Methods.

In this context, we also explored within-individual divergences between homeologous subgenomes within tetraploid assemblies. We found that in all but one case at least 85% of homeologous gene pairs were separated by distances smaller than the relaxed threshold (0.032 substitutions per site) used for species delineation (Fig. 4), consistent with autotetraploidy. The 15% of more divergent homeologous gene pairs were scattered across contigs in the assembly, suggesting they are isolated cases of increased divergence (Oxford dot plots in Supporting Information Section 1). In the remaining case, ilAceEphe1.µ (from host *Acentria ephemerella* [Lepidoptera]), over 40% of homeologous gene pairs exceeded the same-species divergence threshold (Fig. 4A) and many of these divergent homeologues were segregated on separate contigs (Fig. 4B). We note that while we are unable to distinguish between gene copies arising from polyploidisation events *versus* other segmental duplication events for any genome, this high proportion of divergent homeologues in ilAceEphe1.µ may indicate it is the product of a recent hybridisation event between two related diploid individuals (i.e. an allotetraploid). Furthermore, this pattern in ilAceEphe1.µ is unlikely to be the result of a mixed infection of two diploid individuals as the read depths of the four subgenomes are congruent with them belonging to one single genome (Supporting Information Section 1).

**Fig. 4:**
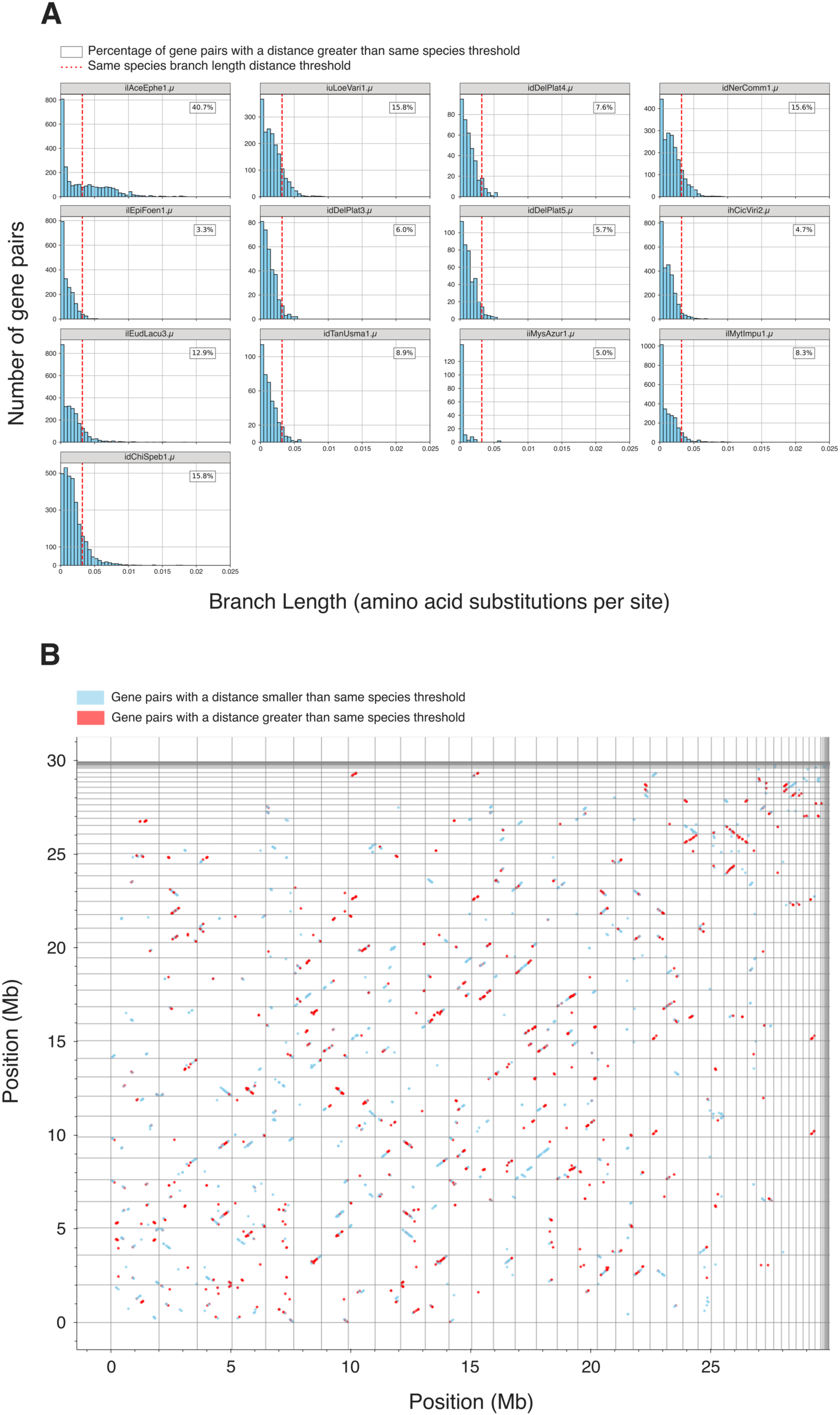
Pairwise phylogenetic branch lengths between homeologous gene pairs in tetraploid genomes. (A) Histograms showing phylogenetic branch lengths (in amino acid substitutions per site) between homeologous gene pairs for thirteen tetraploid genomes. The relaxed branch length threshold for species delineation is highlighted in a dashed red line (0.032 amino acid substitutions per site). The percentage of gene pairs that exceed this same-species threshold is given in a box in the top right of each plot. (B) Oxford dot plot of tetraploid ilAceEphe1.µ (from host *Acentria ephemerella* [Lepidoptera]) using BUSCO genes. Contig boundaries are marked by grey lines. Gene pairs that are less divergent than the same species threshold are in sky blue, while gene pairs that are more divergent than the same species threshold are in red.

### Homeologous genomes coalesce more recently than do species in most tetraploid Microsporidia

With the discovery of tetraploidy in many microsporidian lineages [46–48], one of the key questions is whether tetraploidy is an ancient event shared between multiple (or all) tetraploid microsporidian lineages and subsequently lost by diploid lineages nested within ancestrally tetraploid clades, or is phylogenetically recent and the result of multiple, independent lineage-specific events. The generation of resolved, tetraploid genome assemblies allows us to address this question. In the absence of recombination, gene conversion and sexual reproduction, under ancient tetraploidy we should find that the two homeologous genomes within a tetraploid species have a coalescence deeper than that of the homologous genomes compared between related species [80]. However, if recombination among homeologous genomes occurs, the homeologous genomes within an individual may on average coalesce more recently than they do between species, even while some loci retain a signal of the deep coalescence of the lineages that contributed to the tetraploid. On the other hand, if tetraploidy is the product of recent, independent events, haplotypes will coalesce more recently than species, even in the absence of recombination, and there will be no deep coalescence signal.

We performed the Approximately Unbiased statistical test [81] on multi-copy gene phylogenies for all pairwise combinations of tetraploid microsporidian genomes. We found that the haplotypes in each tetraploid were more similar to each other than they were to genomes from other species (i.e. haplotypes coalesce more recently than species) (Fig. 5). In three species where we had multiple high-contiguity assemblies, the homeologous genomes had coalescences deeper than the within-species coalescence of homologues, implying a single origin of tetraploidy at the base of the species. Thus, the three *Vairimorpha cerenae* genomes, the two genomes assessed in gmOTU1 and the four genomes assessed in gmOTU2 had tetraploid origin coalescences deeper than the within-species coalescence (Fig. 5). Interestingly, the three *Anncaliia algerae* genomes appear more distinct, suggesting reduced flow between the sampled individuals (Fig. 5). As noted above, ilAceEphe1.µ may have arisen from a hybridisation event (Fig. 4). However its diploid ancestors were likely more closely related to each other than they were to other tetraploid genomes sampled in our study, as we observed nearly all coalesces between the homeologues within the tetraploid occurring more recently than they do with other genomes (Fig. 5).

**Fig. 5:**
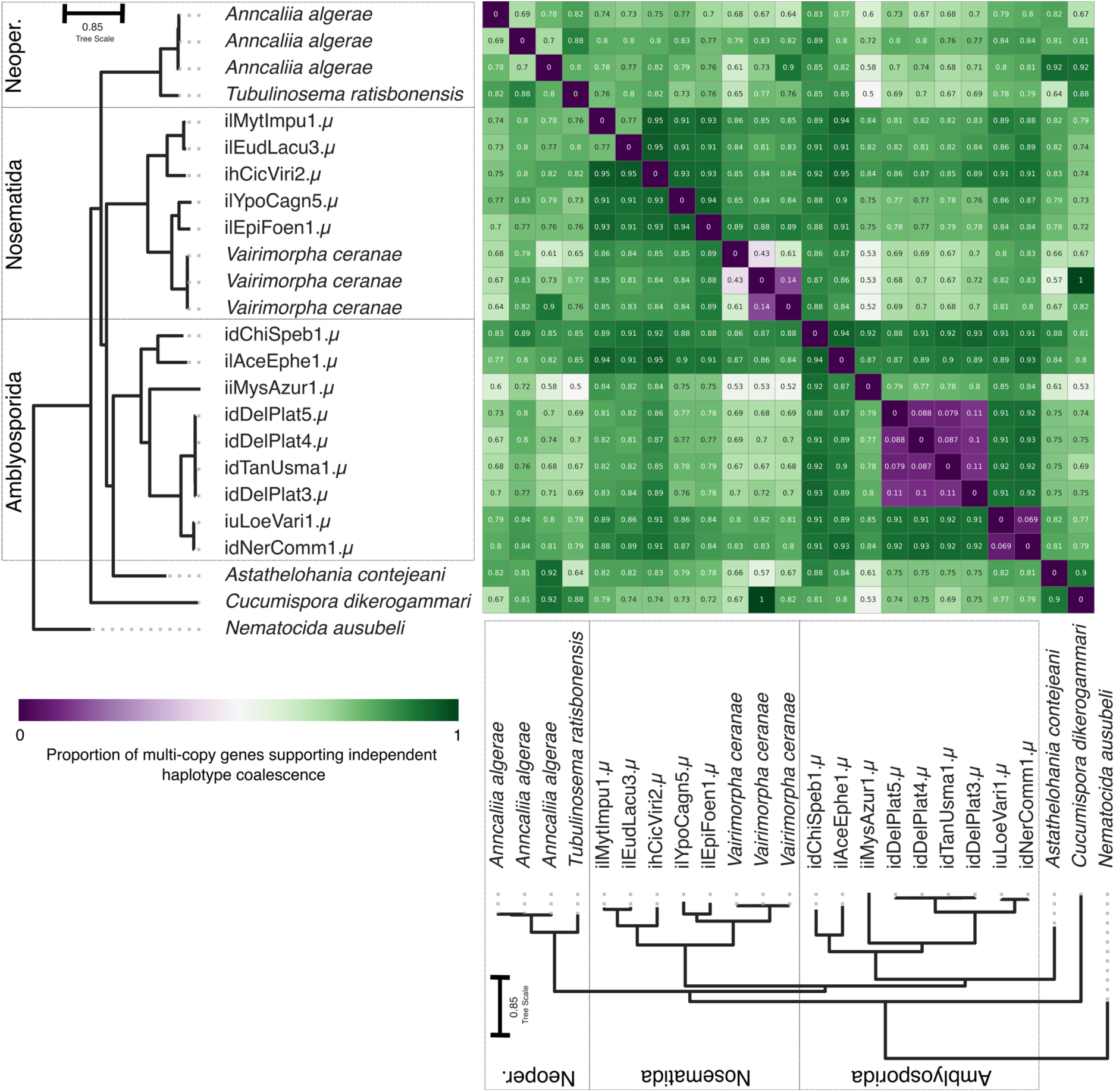
Proportion of multi-copy genes which coalesce prior to genomes. Heatmap showing the fraction of genes that support a more recent homeologue coalescence than between-species coalescence. Fractions greater than 50% are indicated in green, whereas fractions lower than 50% are indicated in purple. The phylogeny is an ASTRAL [69] phylogeny summarising individual phylogenies of 600 BUSCO genes (microsporidia_odb10) [70] from all publicly available tetraploid assemblies and the tetraploid assemblies generated in this study. The branch lengths were estimated using a concatenated alignment of the individual BUSCOs used, with IQ-TREE [71]. The phylogeny is congruent with the phylogeny in Fig. 2.

Because all tetraploid microsporidian genome homeologous subgenomes coalesce more recently than species we cannot distinguish between ancient tetraploidy in a system where recombination, gene conversion and/or sexual reproduction homogenise the two subgenomes; and recent tetraploidy in a system that may or may not undergo recombination and/or sexual reproduction.

### Microsporidian genomes may carry signals of recent recombination

While our coalescence analyses above (Fig. 5) and previous studies suggest that Microsporidia may undergo sexual reproduction [21, 24, 29–40, 42–45], unequivocal signals of recombination have yet to be observed genomically. Phased chromosome-level genomes with known ploidies can help address this by identifying runs of homozygosity between otherwise differentiated homologous chromosomes that may have arisen by recent sexual or non-sexual (i.e. gene conversion) recombination.

For a purged tetraploid genome where all four copies of the genome were reconstructed (iuLoeVari1.µ from host *Loensia variegata* [Psocodea]), we assessed the nucleotide identity patterns between the homologous copies of the largest chromosome. It is striking that this analysis did not identify two pairs of diverged homeologues, but instead suggested a mosaic pattern of pairwise similarity (Fig. 6). In line with this, we found nearly 20% of the tetraploid genome collapsed in the genome assembly (estimated haploid size is ∼17 Mb, whereas tetraploid assembly span is 54.7 Mb). The collapsed regions identified are at the ends of chromosomes, where recombination rates are higher in many organisms [82,83]. Similar patterns were also observed in other high-contiguity genomes (Supporting Information Section 1). These observations are consistent with signatures of recent recombination.

**Fig. 6:**
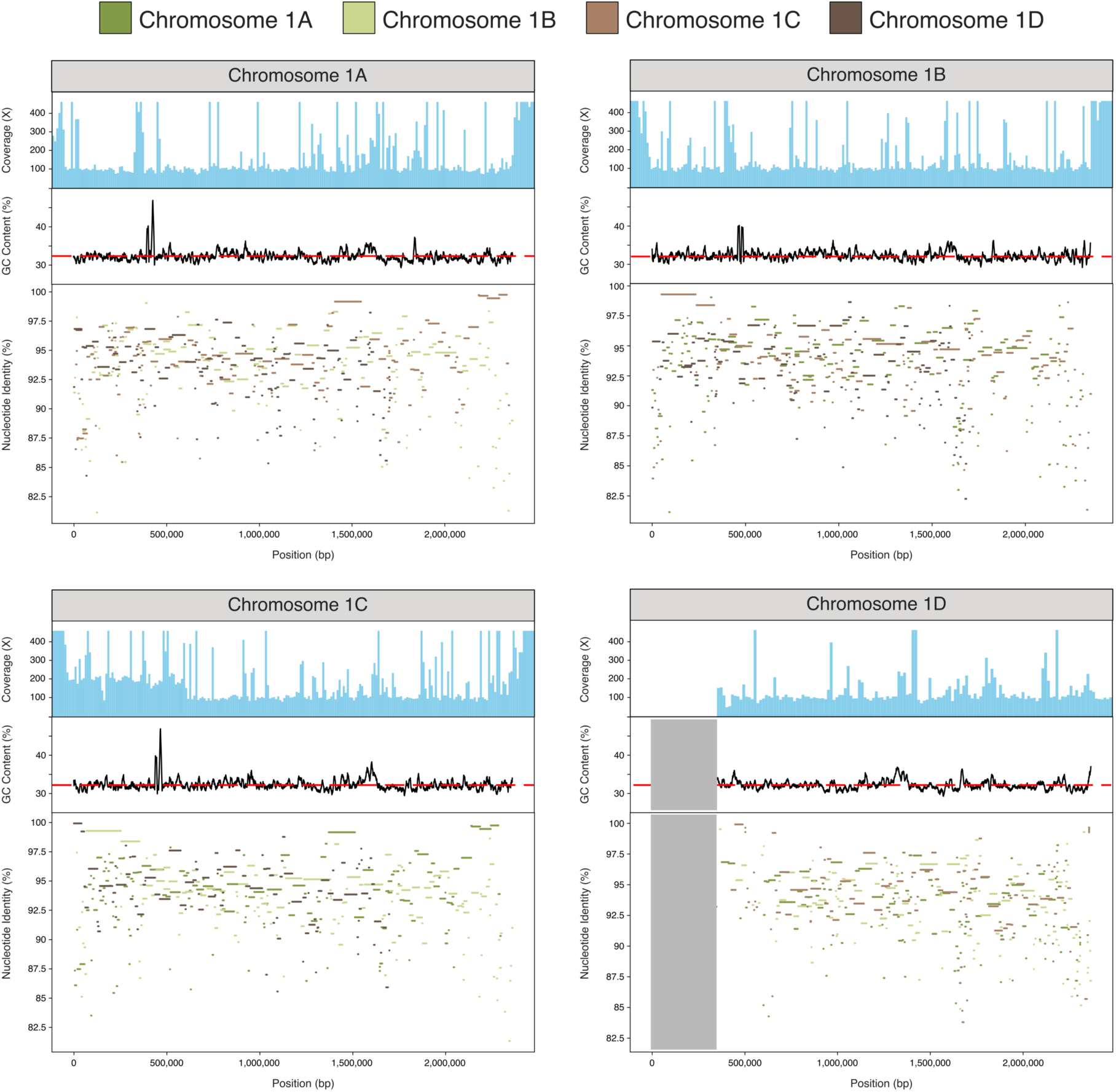
Nucleotide identity (%) between tetraploid iuLoeVari1.µ haplotypes. Each haplotype was compared to the other three haplotypes using minimap2 [84], and nucleotide identity (%) between them was plotted for each reference. Each haplotype has a mosaic pattern of identity to the others. The grey shaded area represents a “missing” segment of chromosome 1D, which we suggest is identical to and thus coassembled as the corresponding portion of chromosome 1C, which has double the expected coverage. The top panel of each plot shows mapped read coverage, and the middle panel displays GC content along the chromosome, with average GC content marked by a dashed red line.

### The tetraploid microsporidian cell contains two diploid compartments

Given the diversity of nuclear conditions in Microsporidia, we sought to understand 3D genome architecture in tetraploids, to determine whether homeologous subgenomes reside in distinct compartments (such as the nuclei of diplokaryons). Thus, we leveraged the availability of high coverage Hi-C data for one of our tetraploid microsporidian genomes, iuLoeVari1.µ from host *Loensia variegata* [Psocodea], to assess physical interactions (chromatin proximities) between the four subgenomes inside the microsporidian cell.

The Hi-C data indicated that for each chromosome, the four homeologous copies have a “two plus two” association (Fig. 7). In addition, looking between chromosomes, these pairs are in turn more likely to interact with other pairs, and the whole genome can be partitioned into two diploid compartments, each containing 20 chromosomes (Fig. 7). Furthermore, the signals consistent with recombination identified above (Fig. 6) occur between the homeologous copies of chromosome 1 within and between these compartments. Taken together, our results suggest that the microsporidian tetraploid genome occurs in two recombining diploid compartments.

**Fig. 7:**
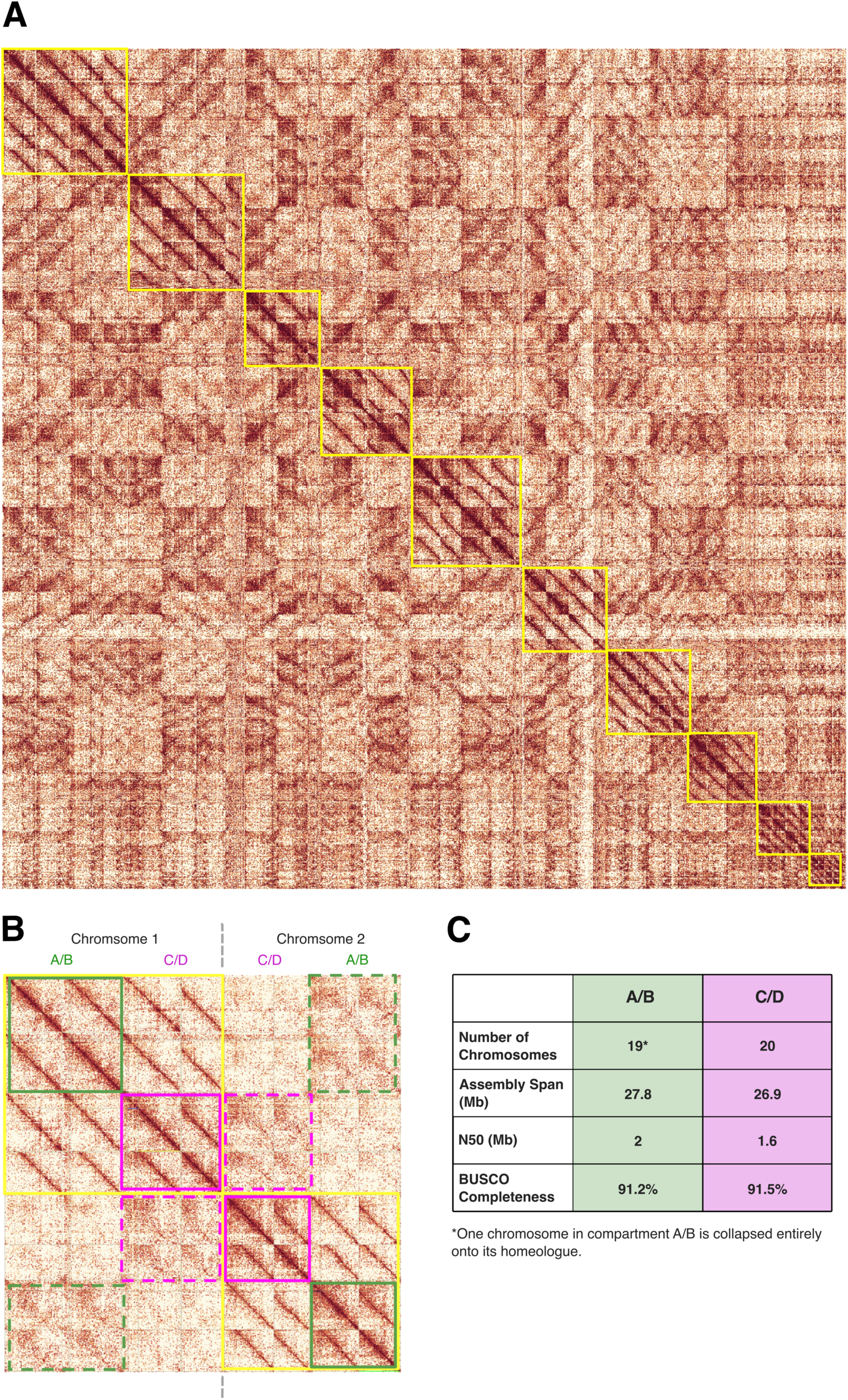
Hi-C heatmap for the tetraploid genome of iuLoeVari1.µ. (A) Hi-C contact map of the tetraploid iuLoeVari1.µ genome (host *Loensia variegata* [Psocodea]). Each chromosome, with its four copies, is highlighted by a yellow box. (B) Hi-C contact map showing the interactions amongst the four copies of chromosome 1 and the four copies of chromosome 2. Green lines highlight interactions belonging to compartment 1, and purple lines highlight interactions belonging to compartment 2. Dotted lines indicate interactions between chromosomes 1 and 2. (C) Summary metrics for the genome assemblies of compartments A/B and C/D. Maps were generated using PretextView [85].

### No evidence of recent rediploidisation in Microsporidia

If tetraploidy was ancestral to Microsporidia or to major lineages within the group, the diploid lineages derived from tetraploid ancestry should show evidence of rediploidisation. Rediploidisation could be achieved by reestablishment of a diploid karyotype by selective loss of one set of chromosomes or through meiotic reduction division without subsequent fertilisation. Alternatively, diploidy could be restored piecemeal by loss or subfunctionalisation of the homeologous copies of each gene [86–90]. Piecemeal rediploidisation should leave a signal in the age of retained homeologous gene pairs, reflecting the divergence between the parental genomes, which would appear as paralogues in the diploidised genome.

We explored age distribution of likely paralogous gene pairs (homologous genes originating from a gene duplication event, called using wgd [86]) on the diploid microsporidian genomes. Relative age was estimated as the synonymous divergence (Ks) between the gene pairs. Piecemeal rediploidisation should be evident as a peak of gene duplicates of the same age [86–90]. Restoration of diploidy through reductive division would not be expected to leave behind such a signal, as the remaining paralogous gene pairs would have had independent origins. We found no peaks of shared divergence in the diploid genomes (Fig. 8). Thus, this suggests that tetraploidy in Microsporidia may result either from recent, independent polyploidisation events; or from an ancestral tetraploid state followed by rediploidisation in diploid lineages through a process similar to that of reductive division in gametogenesis or by inheriting a single nucleus of a diplokaryon.

**Fig. 8:**
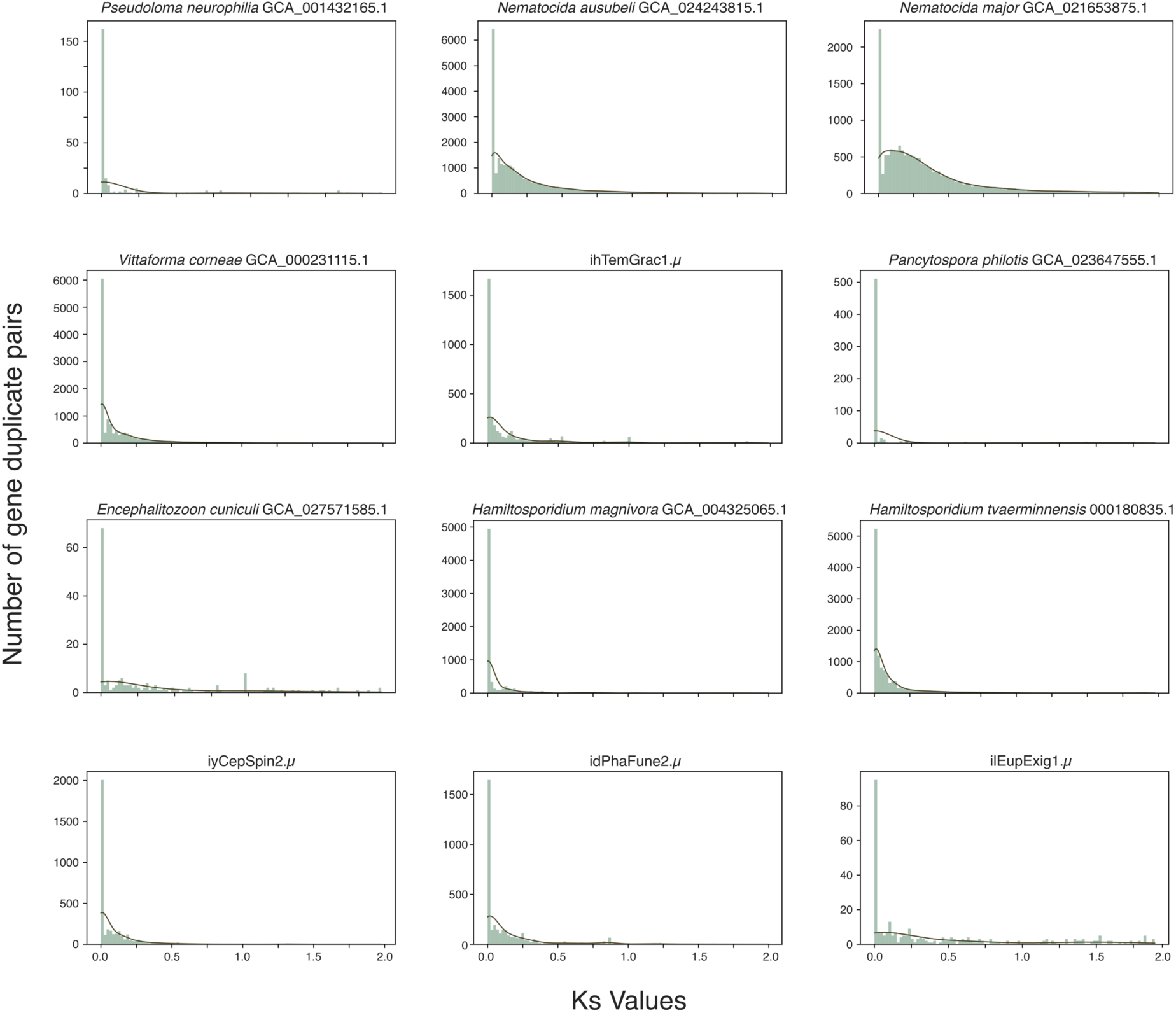
Age distributions of duplicate gene pairs. Histograms showing synonymous divergence (Ks) distributions for candidate paralogous gene pairs from representative diploid genomes from all the major microsporidian groups. No evidence of recent rediploidisation events is seen, as there are no peaks against a background exponentially-decaying distribution coming from small-scale gene duplication events. The y-axis is highly variable due to different gene family expansions occurring in different lineages, yielding larger counts of possible paralogous gene pairs. wgd was used to identify paralogous genes in every genome and compute Ks values [86].

### Segmental duplications are common in Microsporidia

Some microsporidian genomes had large segments differing in copy number from the majority of the genome. These occurred both within and between chromosomes. The duplications varied in length, and preferentially occurred towards the ends of contigs or chromosomes (Self alignment plots in Supporting Information Section 1). For instance, both iyCepSpin2.µ (host *Cephus spinipes* [Hymenoptera]) and idPhaFune2.µ (host *Phania funesta* [Diptera]) have mostly diploid genomes, but carry a level of duplication that generated an identifiable “tetraploid” signal in their k-mer spectra (Fig. 9). Similarly, the k-mer spectrum of iiMysAzur1.µ (host *Mystacides azureus* [Trichoptera]) can be interpreted as either a highly homozygous tetraploid where large segmental duplications have occurred in all the four copies leading to a detectable octoploid signal, or an octoploid genome composed of two distinct tetraploids (Fig. 9). Such cases are common, with some level of segmental duplication observed in nearly all of the 14 polyploid genomes (GenomeScope2 plots in Supporting Information Section 1), but no other phylogenetic or host metadata unites those genomes.

**Fig. 9:**
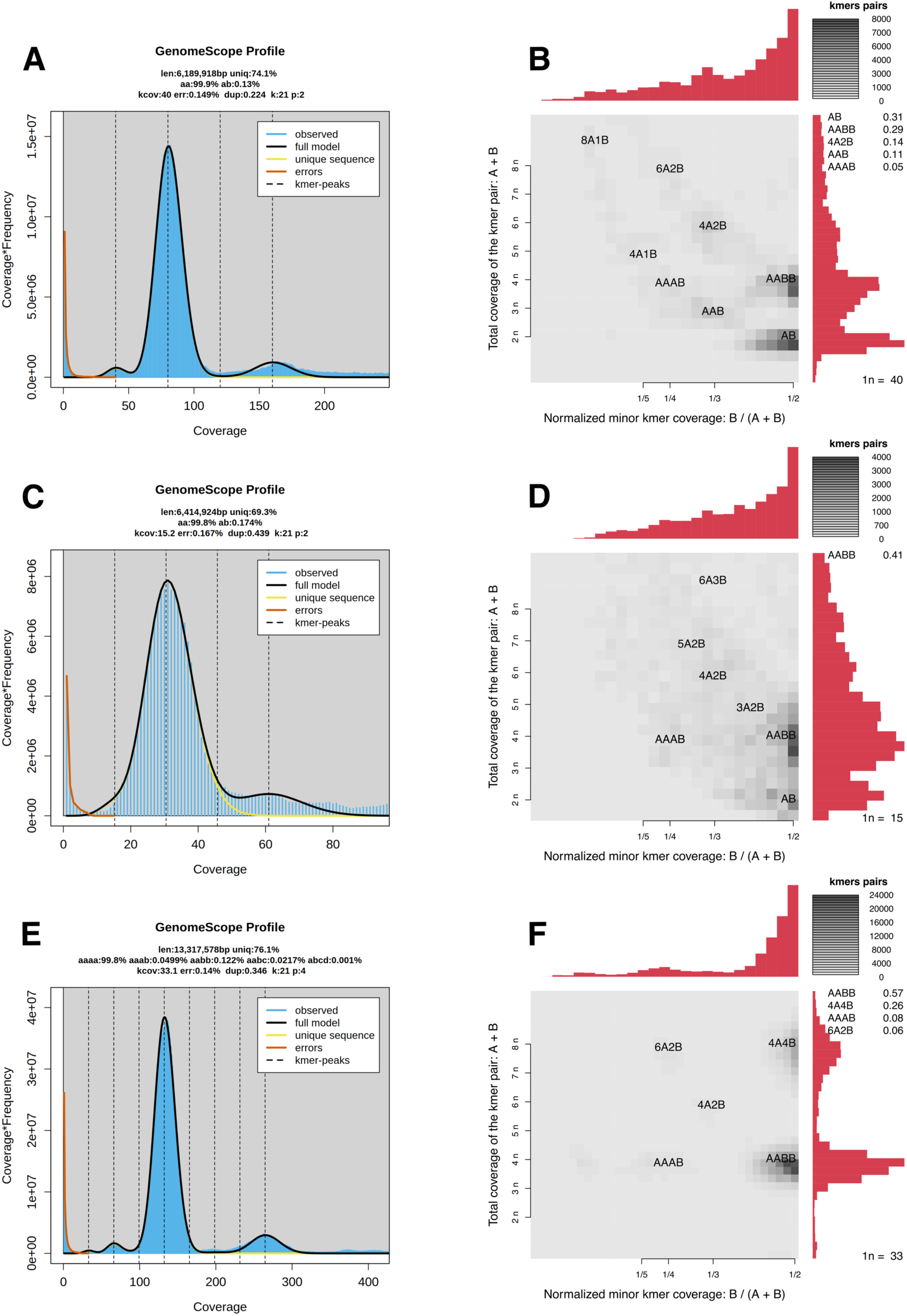
Ploidy inference examples for three microsporidian genomes. GenomeScope2 transformed linear plot and Smudgeplot [91] respectively for (A), (B) diploid iyCepSpine2.µ (host *Cephus spinipes* [Hymenoptera]); (C), (D) diploid idPhaFune2.µ (host *Phania funesta* [Diptera]); and (E), (F) polyploid (tetraploid or octoploid) iiMysAzur1.µ (host *Mystacides azureus* [Trichoptera]). Jellyfish was used to generate the initial k-mer spectra, with k of size 21 [92].

### Evidence for between-homeologue rearrangement in some microsporidian genomes

In addition to segmental duplications, we observe that some tetraploid microsporidian genomes carry signals of between-homeologue rearrangements. Those include inversions, fusions, fissions, and translocations that generated uneven haploid genomes that differed in gene content. To quantify how pronounced this phenomenon is in our genomes, we used a greedy algorithm to bin each tetraploid genome into four subgenome bins. Briefly, the algorithm iterates through contigs from largest to smallest, and appends a contig to a haplotype if the duplication in that contig does not exceed a specified threshold (see Github: https://github.com/Amjad-Khalaf/gerbil for implementation).

idChiSpeb1.µ (host *Chironomus sp.* [Diptera]) is a perfect tetraploid, with no assembly collapse, and thus can be binned into four equal subgenomes for the vast majority of duplication thresholds tested (Fig. 10). Similarly, we were able to partition the genomes of iuLoeVari1.µ (host *Loensia variegata* [Psocodea])) and idNerComm1.µ (host *Neria commutata* [Diptera)] recovering two incomplete subgenomes showing some genome collapse (Fig. 10). On the other hand, for three other genomes including ilAceEphe1.µ (host *Acentria ephemerella* [Lepidoptera]), ilMytImpu1.µ (host *Mythimna impura* [Lepidoptera]), and ihCicViri2.µ (host *Cicadella viridis* [Hemiptera]) we were unable to recover a haplotype with completeness similar to that of the unbinned genome assembly without also retaining a higher amount of duplication than expected (Fig. 10, Supporting Information Section 4). In line with this, the self-alignment dot plots of those genomes show numerous rearrangements and translocations (Self alignment plots in Supporting Information Section 1). We also found evidence of similar rearrangements and unevenness in other diploid and tetraploid purged microsporidian genomes, and in some genomes without known ploidies (Self alignment plots in Supporting Information Section 1). Together, these results provide evidence for extensive between-haploid subgenome rearrangement in microsporidian genomes, resulting in uneven gene distribution between haploid subgenomes. We examined whether between-haplotype rearrangements were associated with any genome meta-data, and found no phylogenetic, host, or ecological association linking them.

**Fig. 10:**
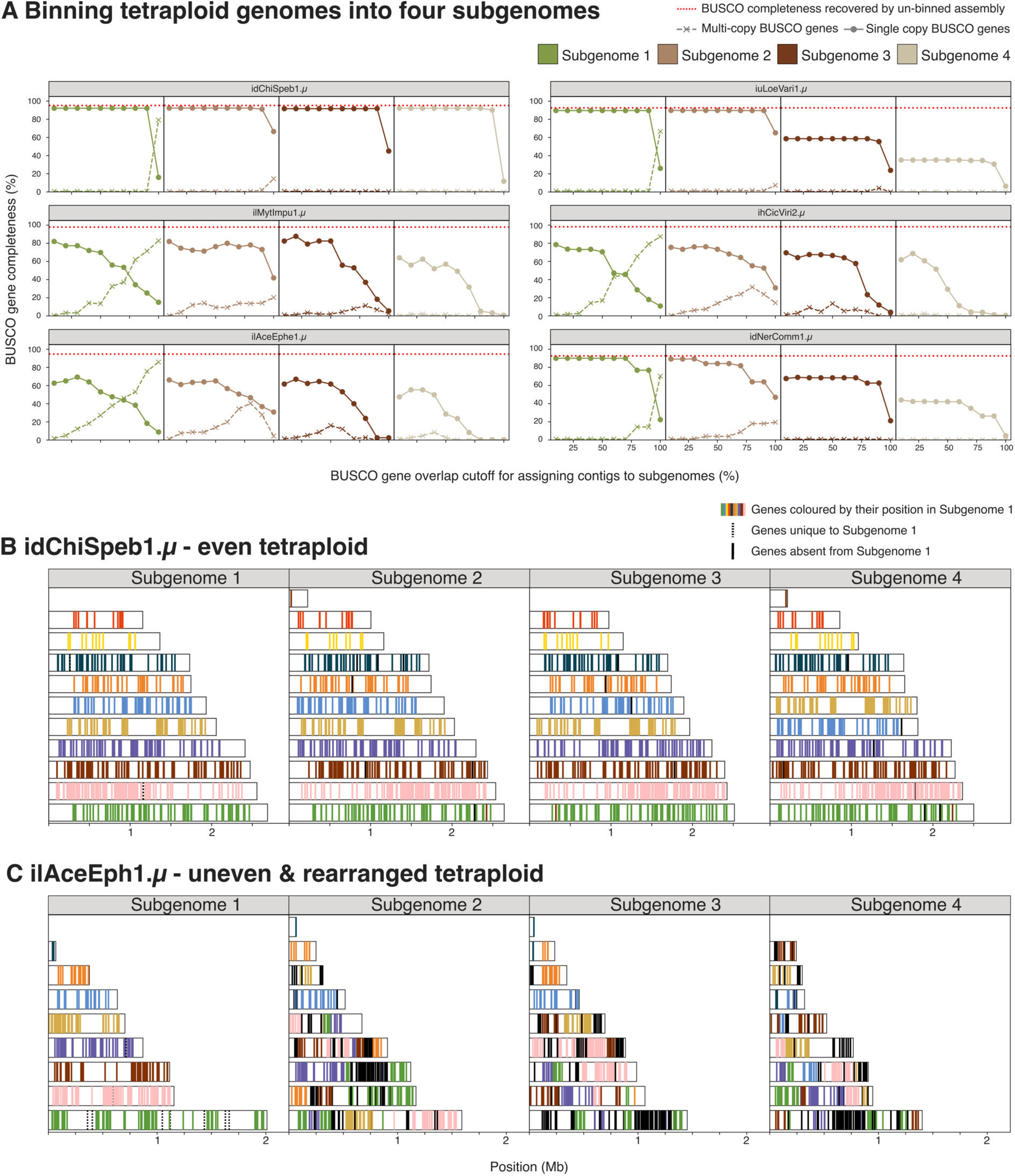
Binning tetraploid genomes into four subgenomes using BUSCO genes. (A) Using a greedy algorithm, we iterated through contigs from largest to smallest, appending a contig to a haplotypic subgenome if the duplication in that contig does not exceed a specified threshold (x axes in the figure). Single copy BUSCO gene completeness is marked by circles and multi-copy BUSCO gene completeness is marked by crosses. A red dashed line denotes the BUSCO completeness score of the unbinned assembly. For (B) idChiSpeb1.µ and (C) ilAceEphe1.µ, we plotted the largest 10 contigs in subgenome 1 with their BUSCO genes, and coloured these genes in the other subgenomes by their positions in subgenome 1.

### Large-scale chromosomal rearrangements between microsporidian groups

Given the above observations of synteny breakage between subgenomes within tetraploids, we expect to observe fractured synteny between species. We mapped the relative position of conserved orthologous BUSCO genes in all chromosome-level genome assemblies. Confident reconstruction of ancestral linkage groups for all Microsporidia was not possible, likely because the rearrangement rate was indeed too high (see Supporting Information Section 5 for details).

Comparing closely-related genomes showed a pattern of dynamic change of microsporidian linkage groups. The *Encephalitozoon* species genomes were highly syntenic, with the exception of rearrangement involving a single linkage group [61] (Fig. 11 A). ilEupExig1.µ (host *Eupithecia exiguata* [Lepidoptera]), placed basal to *Encephalitozoon* species, had six chromosomes that could be derived through either five pairwise fusions of the 11 chromosomes of *Encephalitozoon* species, or an ancestral karyotype that was subject to fission in *Encephalitozoon*. *Vairimorpha necatrix* was sister to the *Encephalitozoon-* ilEupExig1.µ clade, and had 11 chromosomes, but these did not correspond to the 11 found in *Encephalitozoon* species and did not simply confirm the karyotype of ilEupExig1.µ as being ancestral or derived. The karyotype of the enterocytozoonid iyOphElle1.µ, sister to the nosematids (*V. necatrix, Encephalitozoon* and ilEupExig1.µ), had 12 chromosomes, the largest of which is syntenic with the largest chromosome of *V. necatrix*, and thus suggests that the splitting of this chromosome in *Encephalitozoon* and ilEupExig1.µ is derived. This large linkage group was also found in the neoperezeiid *Antonospora locustae*, sister to the Encephalitozoonida, in the orphan lineage species *Hamiltosporidium tvaerminnensis* and in two genomes on Ambylosporidia species, suggesting it was likely present in the last ancestor of all microsporidia analysed (Supporting Information Section 5).

**Fig. 11:**
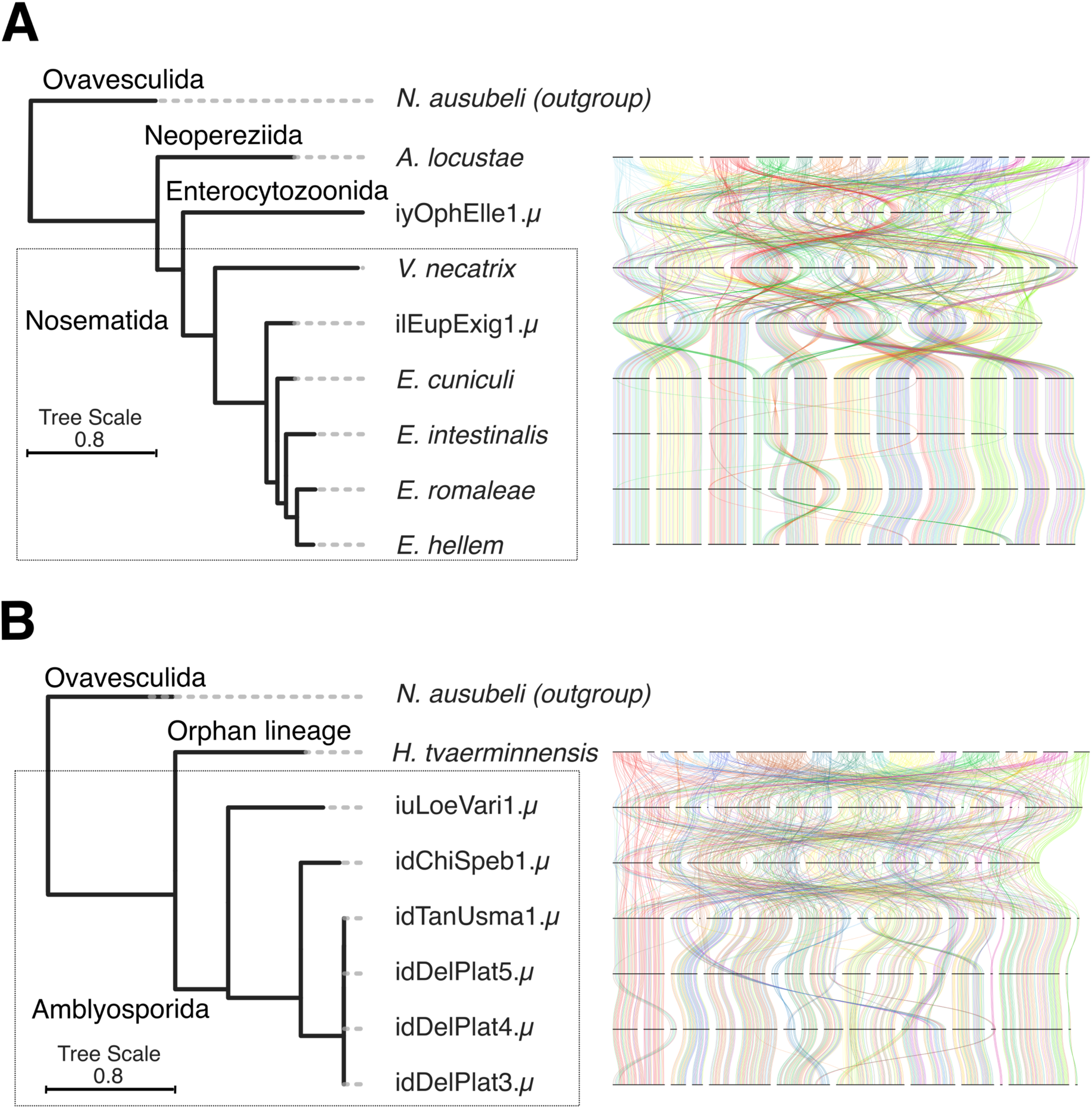
Synteny plots of chromosomal microsporidian genome assemblies. Genome-wide synteny plots of all chromosomal microsporidian genome assemblies for (A) Enterocytozoonida, Nosematida, and Neopereziida; and (B) Amblyosporida and the Orphan lineage. Each line represents a single-copy BUSCO (microsporidia_odb10) [70]. In (A) BUSCOs are painted by their chromosomal position in *A. locustae*, while in (B) they are painted by their chromosomal position in *H. tvaerminnensis*. Plots were generated by modifying the ribbon plotting script in https://github.com/conchoecia/odp [96]. The attached phylogeny is an ASTRAL phylogeny summarising individual phylogenies of 600 BUSCO genes (microsporidia_odb10) [70]. The branch lengths were subsequently estimated using a concatenated alignment of the individual BUSCOs used, with IQ-TREE [71].

The orphan lineage-Ambylosporidia group showed a similar general pattern of linkage group conservation between close relatives and members of the same OTU, coupled with major rearrangements between clades (Fig. 11 B). We were unable to infer a robust set of putative ancestral linkage groups for these genomes using syngraph [93] or unsupervised clustering of loci based on their chromosomal occupancy [94,95], likely because of with the high frequency of rearrangements observed (see Supporting Information Section 5).

## Discussion

### Forty new genome assemblies, a revised phylogeny, and the future of species delineation in Microsporidia

In this study, we present 32 new complete microsporidian genome sequences (including eight chromosome-level genomes, seven of which, to our knowledge, were scaffolded with the first implementation of Hi-C data on Microsporidia), and eight additional partial genome sequences (with BUSCO completeness score < 70%). While not being the target organisms in the sequencing efforts used to generate these genome assemblies, most of the assemblies presented in this study have comparable or higher contiguity and BUSCO completeness than published microsporidian genomes (Fig. 2). Our genome assembly process provides an accessible and successful approach for the assembly of microsporidian genomes from their host sequencing data, and can be translated to other cobionts where a wealth of sequenced host genomes is available.

The presented microsporidian genomes derive from five of the seven major microsporidian clades as defined by Bojko and colleagues [4]. They allowed us to revise microsporidian phylogeny, notably changing the position of Glugeida to the sister group of the ancestor of the ‘orphan lineage’ and Amblyosporida (Fig. 2).

We show that genomic data support previous species delineations based on morphology, histopathology, and cell culture, and suggest divergence thresholds that differentiate species. While morphological and histopathological data remain important for identifying Microsporidia to species, it is likely that serendipitous discovery through large-scale genomic sequencing will be a major source of new microsporidian isolate genomes in the future. In prokaryotes, genomic metrics for species delineation, such as Average Nucleotide Identity (ANI) [97], are well accepted. The application of similar metrics to taxon delimitation in eukaryotes is in its infancy [98].

### Tetraploidy in Microsporidia

From theoretical and observational grounds, the long-term maintenance of tetraploidy is unlikely. Usually, a new tetraploid lineage will revert to effective diploidy, either by losing one set of chromosomes or by reestablishment of exclusive pairing between homologues and doubling of the diploid chromosome number. The reestablishment of diploidy (rediploidisation) leaves a signature of whole genome duplication (WGD) in descendants, involving loss of copies of many genes and subfunctionalisation of retained genes [99]. WGD has been common during evolution of plants, animals and fungi, and has been proposed to be a significant driver of organismal complexity and adaptive evolution [50]. The situation in Microsporidia differs from classical yeast, flowering plant and animal models of tetraploidy and rediploidisation, as the constituent genomes appear to be largely maintained intact and show signatures of ongoing recombination between all four genomes.

In some microsporidian genomes, we observed nearly perfect synteny between haplotypes. However, we observed a striking level of between-haplotype rearrangement in three microsporidian genomes, two autotetraploids (ilMytImpu1.µ and ihCicViri2.µ) (Fig. 10), and a genome with circumstantial evidence of being the result of a recent hybridisation event between two closely related diploid genomes (Fig. 4, Fig. 10). We also see similar signatures, albeit on smaller scales, in other genomes (Self alignment plots in Supporting Information Section 1). In other taxa, such rearrangement is often associated with hybrid origin in polyploids, for example in synthetic plant tetraploids such as *Brassica* allotetraploids [100–104] and autotetraploid *Arabidopsis thaliana* [105], and in the multiple origins of tetraploidy in lager brewing yeast *Saccharomyces pastorianus* [106–108]. Such rearrangement may be a common genomic response to the “genome shock” of the origin of tetraploidy [109,110].

We cannot currently distinguish a general genomic shock hypothesis for the origin of these rearranged microsporidian genomes from other possibilities, such as independent genetic disruption of reproductive processes, as seen in a number of yeast isolates with similar karyotype patterns [111]. Population-level whole-genome sequencing of selected species, over time (both in nature and in lab cell cultures), is required to better define this phenomenon and identify its root causes.

### A diploid/tetraploid mating system as a potential reproductive model for Microsporidia

Understanding the origin of tetraploidy and its biology in Microsporidia has significant implications. If tetraploidy is ancient, Microsporidia would represent a rare case of a stable, species-rich tetraploid lineage, offering a unique opportunity to study genome evolution under these conditions. On the other hand, if tetraploidy is recent, Microsporidia would represent a clade with an unusually high propensity for polyploidisation. Additionally, if ploidy influences host specificity, this could inform strategies to mitigate the impact of Microsporidia on aquaculture and beekeeping or their potential as biological control agents, such as for malaria. Furthermore, a deeper understanding of their reproductive behaviour could also aid in both managing and leveraging these organisms.

We identified many tetraploid microsporidian genomes, the majority of which are likely autotetraploids (Fig. 4), but no concrete evidence that tetraploidy was the ancestral state in Microsporidia. We observed that homeologous subgenomes coalesce more recently than do the genomes of different species in all cases (Fig. 5). However, this does not allow us to distinguish between ancient shared tetraploidy versus recent independent tetraploidy, as both models are likely to be indistinguishable in the presence of recombination within and between homeologous subgenomes during sexual reproduction. Similarly, in diploid lineages nested within tetraploids, we did not observe signals congruent with classic piecemeal rediploidisation (Fig. 8). This suggests that tetraploidy in Microsporidia is independently acquired in the polyploid clades, or that diploid lineages have restored diploidy through one-step mechanisms that eliminate one diploid set of chromosomes. This could arise if a lineage was founded from one diploid compartment (Fig. 7), or through meiotic reduction division without subsequent fertilisation.

If tetraploidy is the result of recent, independent events, a minimum of 15 events is predicted. We thus suggest that it is more likely that a propensity for tetraploidy is ancient in Microsporidia, and that the diploid lineages nested within tetraploids represent isolates or lineages that have undergone reductive division.

One of our key findings is that tetraploid microsporidian genomes are likely organised into two diploid compartments (Fig. 7), with evidence of recombination both within and between compartments (Fig. 6). Taking all our findings together, we propose that the compartments are the two nuclei of a diplokaryon (i.e. each nucleus is diploid), that Microsporidia undergo occasional sexual reproduction in a process that mirrors fungal reproduction (Fig. 12), and that chromosomes independently reassort into the compartments in different individuals. There is extensive morphological evidence of plasmogamy (cell fusion) and karyogamy (nuclear fusion) in Microsporidia [21,24,29–40,42,43], and these processes may reflect this proposed sexual cycle.

**Fig. 12:**
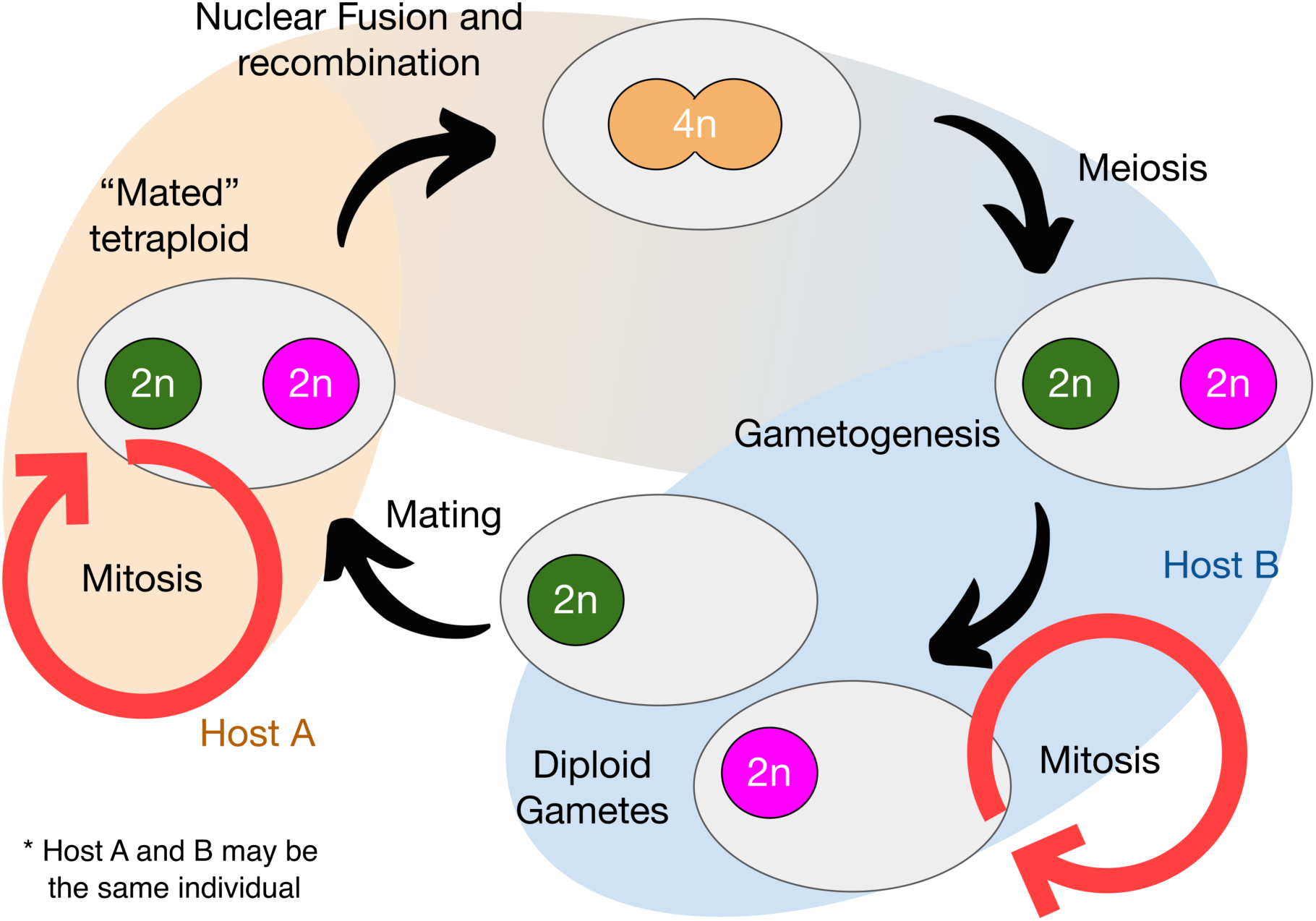
Proposed generalised lifecycle for Microsporidia. Our proposed model posits that each nucleus is a diploid, and that microsporidian reproduction mirrors reproduction in Fungi with stages similar to karyogamy, plasmogamy, and a stable “heterokaryon” (known as a diplokaryon in Microsporidia). Importantly both the diplokaryotic and monokaryotic phases are parasitic, and species may spend most of their lifecycle in one or the other phase, giving rise to “diploid” and “tetraploid” lineages.

In this model both diploids and tetraploids can function as infective agents (Fig. 12). Support for this also comes from life cycle observations. Several microsporidians exhibit complex life cycles, generating multiple spore types with different infective potentials from a variety of hosts. For example, diplokaryotic *Edhazardia aedis* spores infect adult mosquitoes, producing monokaryotic spores that go on to infect larvae, generating diplokaryotic spores [31]. *Amblyospora connecticus* cycles between mosquito and copepod hosts using monokaryotic and diplokaryotic spores for infecting each host respectively [112]. This alternation of generations with different ploidies could explain why the four genomes in tetraploids remain intact, and have not been subjected to the usual processes of rediploidisation, as random assortment of genomes into the diploid would select for fully functional diploids whichever pair was combined.

In the proposed model each diplokaryon nucleus is diploid, and their mitosis requires only pairing of two homologous chromosomes, as in any diploid. In the proposed tetraploid fusion, however, pairing and correct partitioning of the chromosomes to diploid daughter nuclei requires that all four homologous chromosomes are recognised and sorted coherently. In diploid cells, homologous chromosome pairing is mediated through sequence recognition. Minimally, the four copies of each chromosome in the proposed tetraploid fusion must carry some signature that means they can be recognised even if they diverge in sequence elsewhere such that they can no longer recombine.

Our model predicts that there should be diploid, “gametic” forms for all the tetraploid Microsporidia, and that tetraploid forms may exist for described diploid lineages. We note that it remains unclear how ploidy maps on to life stages in most species, and this information is crucial to underpin the nature of microsporidian reproduction. Generation of single-cell, whole-genome data for Microsporidia, focussing especially on species with complex life cycles and multiple spore types would be highly informative.

While the diploidy of the microsporidian nucleus is broadly in line with morphological data, with most diploid species being monokaryotic, and most tetraploid species being diplokaryotic [46,47], there are two exceptions. Namely, tetraploid *Agmasoma penaei,* which possesses monokaryotic spores (suggesting each nucleus is tetraploid), and diploid *Vittaforma corneae,* which possesses diplokaryotic spores (suggesting each nucleus is haploid) [46,113]. Given the morphological descriptions are not from the same isolates used to predict their ploidy, it is possible that these species are polymorphic, as observed in Microsporidia with complex, alternating lifecycles and multiple spore types [41,114]. Future sampling of those lineages in particular will be crucial to whether each nucleus is truly diploid in Microsporidia.

In this work, we generated 40 high-quality genome assemblies, found signals consistent with recombination, and provided evidence suggesting that microsporidians are likely autotetraploids. While the timing and number of polyploidisation events remains uncertain, we propose that tetraploidy is an ancient feature of Microsporidia, with diploid lineages representing “reduced” forms. Population-level whole-genome sequencing, combined with longitudinal imaging in nature and laboratory cultures, will be crucial to illuminate the true nature of polyploidy, and its relationship to the lifecycle in Microsporidia.

## Materials and Methods

### Data

Samples sequenced as part of the DToL project [62] were processed by the Tree of Life core laboratory and the Scientific Operations core at the Wellcome Sanger Institute. This process typically relies on using different parts of one individual, or two closely related individuals, to generate long-read DNA sequencing data and Hi-C short-read sequencing data. Because of this, specimens identified as microsporidian-infected from long-read sequencing data did not often have Hi-C data available.

To ensure we were able to generate Hi-C data for a subset of the microsporidian genomes we explored, we sampled 650 individual flying insects on the Wellcome Genome Campus using a Malaise trap in the summer of 2023. These specimens were first bisected, and DNA for long-read sequencing was extracted from one half. The other half was kept at −80°C for later Hi-C sequencing. We identified infected insects by PCR amplification testing of the long read DNA extracts using a microsporidia-specific amplicon panel [115,116]. Six specimens were identified as microsporidia-positive, and library preparation and genome sequencing of those six individuals, with ToLIDs idDelPlat3.µ, idTanUsma1.µ, idChiSpeb1.µ, idDelPlat4.µ, idLucSpea1.µ, and idDelPlat5.µ, was performed by the Scientific Operations core at the Wellcome Sanger Institute.

BioSpecimen identifiers for each dataset used in this study are listed in the Supporting Information Section 1, along with the microsporidian genome assemblies.

### ToLIDs and OTUs

We use Tree of Life identifiers (ToLIDs) of the host individuals that were sequenced at the Wellcome Sanger Institute, with the suffix “.µ”, to refer to the microsporidian genome assemblies that resulted from them (see https://id.tol.sanger.ac.uk/ for more information). The full ToLIDs the microsporidian genome assemblies are released under are listed in Supporting Information Section 1. Additionally, we use “gm” as a prefix for each OTU, in line with ToLID notation (g for “Fungi”, and m for “Microsporidia”).

### Genome assembly

We identified 34 individual specimens sequenced for DToL as likely to be infected with a microsporidian using a MarkerScan [66] screen of their preliminary genome assemblies (generated using hifiasm version 0.19.9 [117]). For each infected individual, we concatenated the primary and alternate preliminary genome assemblies.

To positively identify microsporidian sequences, BlobToolKit [118] was run on each concatenated preliminary genome assembly. Contigs were filtered (“Filtering Step 1” in Supporting Information Section 1) by a combination of average read coverage, GC content, and taxonomic classification of contigs using upper and lower bounds that retained contigs that had BLASTx matches mostly to microsporidian proteins, and excluded contigs that had a majority of database matches to proteins from to other taxa. The specific filtering parameters used for each genome assembly are reported in Supporting Information Section 1.

The PacBio single-molecule HiFi long reads belonging to the preliminary assembly were aligned to the filtered microsporidian contigs using minimap2 (version 2.28) [84], and aligned reads were isolated using samtools (version 1.19.2, MAPQ = 255) [119]. To assess read coverage and ploidy, a k-mer spectrum was generated for the isolated reads using Jellyfish (k = 21, version 2.2.10) [92] and analysed using GenomeScope2 (version 2.0) [91]. Where samples had high average read coverage and reliable ploidy estimation, Smudgeplot (version 0.4.0 “Arched”) was also run to confirm the ploidy assessment [91].

The isolated PacBio HiFi reads were then reassembled using hifiasm (version 0.19.9, parameter −l0 was used to disable purging) [117]. The contigs from the reassembly were filtered again (using https://github.com/Amjad-Khalaf/BubblePlot, “Filtering Step 2” in Supporting Information Section 1) by average read depth, GC content, and taxonomic classification of contigs, with upper and lower bounds that retained all contigs that contained microsporidian BUSCO proteins (microsporidia_odb10, version 5.4.6) [70], and excluded contigs that contained proteins mostly belonging to other taxa. The specific filtering parameters used for each genome assembly are reported in Supporting Information Section 1.

In cases where ploidy was successfully estimated from the isolated reads, the ploidy status of the cleaned reassembly was assigned, or marked as unresolved (“NA” in Supporting Information Section 1).

If Hi-C data was available for the same individual that the long read data had been generated from, Hi-C reads were mapped to the cleaned reassembly using bwa-mem2 (version 2.2.1) [120], and the resulting alignment files were converted to contact maps using Juicer Tools (version 1.8.9) [121], bedtools (version 2.31.1) [122], and PretextMap (version 0.19) [123]. If Hi-C data provided sufficient signal, contigs were then manually scaffolded using PretextView (version 0.2.5) [85]. A haploid representation of the genome assembly was generated by selecting the most contiguous copy of each chromosome with the least gaps. This was possible for seven microsporidian genome assemblies.

If Hi-C data was not available, but the cleaned reassembly possessed sufficient coverage to estimate ploidy using GenomeScope2 (version 2.0) and Smudgeplot (version 0.4.0 “Arched”) [91], purge_dups was used to generate a haploid representation of the genome assembly [124]. This process was followed for eight microsporidian genome assemblies.

In the cases where Hi-C data was not available, or the cleaned reassembly did not possess sufficient coverage to estimate ploidy, or the cleaned reassembly showed high levels of rearrangements between its subgenomes (regardless of having been assigned a ploidy), purge_dups [124] was not run. For these assemblies, the unresolved cleaned reassembly was considered the final assembly and used in further analyses. This was the case for 27 microsporidian genome assemblies. The data available and the paths followed for each microsporidian genome assembly are outlined in Supporting Information Section 1.

The genome assembly and read metrics for all intermediate steps for each microsporidian genome assembly are reported in Supporting Information Section 1.

### Publicly available microsporidian genome assemblies

On the 1st of January 2025, we downloaded all microsporidian genome assemblies available in the NCBI Genome database. This retrieved 106 genome assemblies, whose accession numbers are listed in Supporting Information Section 1.

### Alignment and phylogeny

Unless stated otherwise, all phylogenies were generated by identifying orthologous proteins using BUSCO (microsporidia_odb10, version 5.4.6.) [70], aligning each locus across all genomes using MAFFT (version 7.525) [125], inferring a tree for each of them using IQ-TREE (version 2.3.4) [71], and summarising all the resulting gene trees into one species tree using ASTRAL (version 5.7.8) [69]. The branch lengths were subsequently estimated using a concatenated alignment of the individual BUSCOs used, with IQ-TREE [71]. The model chosen according to MFP was “Q.yeast.I.G4”. In the case of multi-copy genes, one of the copies was chosen randomly. This was done because the majority of multi-copy genes across all the genomes showed that haplotypes displayed clear monophyly (see Results).

To infer species membership of unidentified genomes, we calculated average pairwise divergence between all possible genome pairs using the BUSCO gene phylogeny, and defined upper bounds for within-species divergence based on the genomes from identified microsporidian species from public data. We also explored generating a same-species threshold for each gene, using the distribution of branch lengths between genomes classified as belonging to the same species. Our results were consistent with those based on the whole-genome phylogeny branch lengths (see Supporting Information Section 3).

### Assessing if haplotypes coalesce prior to genomes

For every possible combination of any two tetraploid genomes from the list of complete genomes generated in this study, and all publicly available genomes, a tree topology test was performed using IQ-TREE (version 2.3.4) [71] on multi-copy BUSCO gene trees. Support for the topology where each genome’s haplotypes displayed clear monophyly was statistically assessed using the Approximately Unbiased Test [71,81]. Unpurged genome assemblies were used where available.

### Inference of historical rediploidisation events

To infer potential historical de-polyploidisation events, wgd (version 2) was run on each genome from the list of complete genomes generated in this study, and all publicly available genomes, to infer paralogous genes and compute their Ks values [86].

### Genome annotation

Each final microsporidian genome assembly was annotated for coding sequences using Prokka (version 1.14.6) [72], and for repeats using RepeatModeler (version 2.0.5) and RepeatMasker (version 4.1.7) [73,74]. Coding sequence annotations, and repeat annotations for each genome assembly are reported in Supporting Information Section 1. While Prokka was designed for prokaryotic genome annotation, we elected to use it on microsporidian genomes due to their extremely low number of introns [126,127], and the fact that no functional annotations were required in this piece of work.

### Between-haplotype rearrangements

To illustrate between-haplotype rearrangements in some of our genomes, we developed a greedy algorithm that bins each of our unpurged tetraploid genomes into four subgenomes based on BUSCO gene completion and duplication (see Github: https://github.com/Amjad-Khalaf/gerbil for implementation). After sorting contigs by from largest to smallest, our algorithm iterates through contigs and assigns a contig to a subgenome bin if the duplication that contig would add to its proposed subgenome bin does not exceed a specified threshold.

The discussed patterns can be highlighted by examining synteny between the recovered haplotypes for one of the duplication thresholds tested (the x-axis in Fig. 10A). For idChiSpeb1.µ and ilAceEphe1.µ, we plotted the largest 10 contigs in haplotype 1 with their BUSCO genes, and coloured these genes in the other haplotypes by their positions in haplotype 1 (Fig. 10). As expected, the haplotypes of idChiSpeb1.µ show nearly perfect synteny with one another. However, ilAceEphe1.µ display a large number of rearrangements and haplotype-unique BUSCO genes (as seen by the pattern in Fig. 10). In line with this, nearly all of idChiSpeb1.µ’s BUSCO genes are in 4 copies, distributed across 4 haplotypes (Supporting Information Section 4). On the other hand, the vast majority of ilAceEphe1.µ’s BUSCO genes are in less than 4 copies (despite its haploid coverage exceeding 48X, as seen in Supporting Information Section 1), and its BUSCO genes are not evenly distributed across its haplotypes. For instance, some BUSCO genes occur in 3 copies, all present in a single haplotype (Supporting Information Section 4). Together, these observations highlight the degree of rearrangement and unevenness in ilAceEphe1.µ, and similar scenarios are seen for ilMytImpu1.µ and ihCicViri2.µ.

## Supporting information

Supporting Information Guide

## Acknowledgements

We thank Dr Lewis Stevens, Dr Jamie Bojko, and Dr Yuliya Y Sokolova for their insight, their kindness, and generosity with their time in discussing these results with us over the last year. We also warmly thank our colleagues at the Tree of Life Programme, Wellcome Sanger Institute for their support, and comradery. This work was funded in whole by the Wellcome Trust (grant number 220540/Z/20/A).

## Author contributions

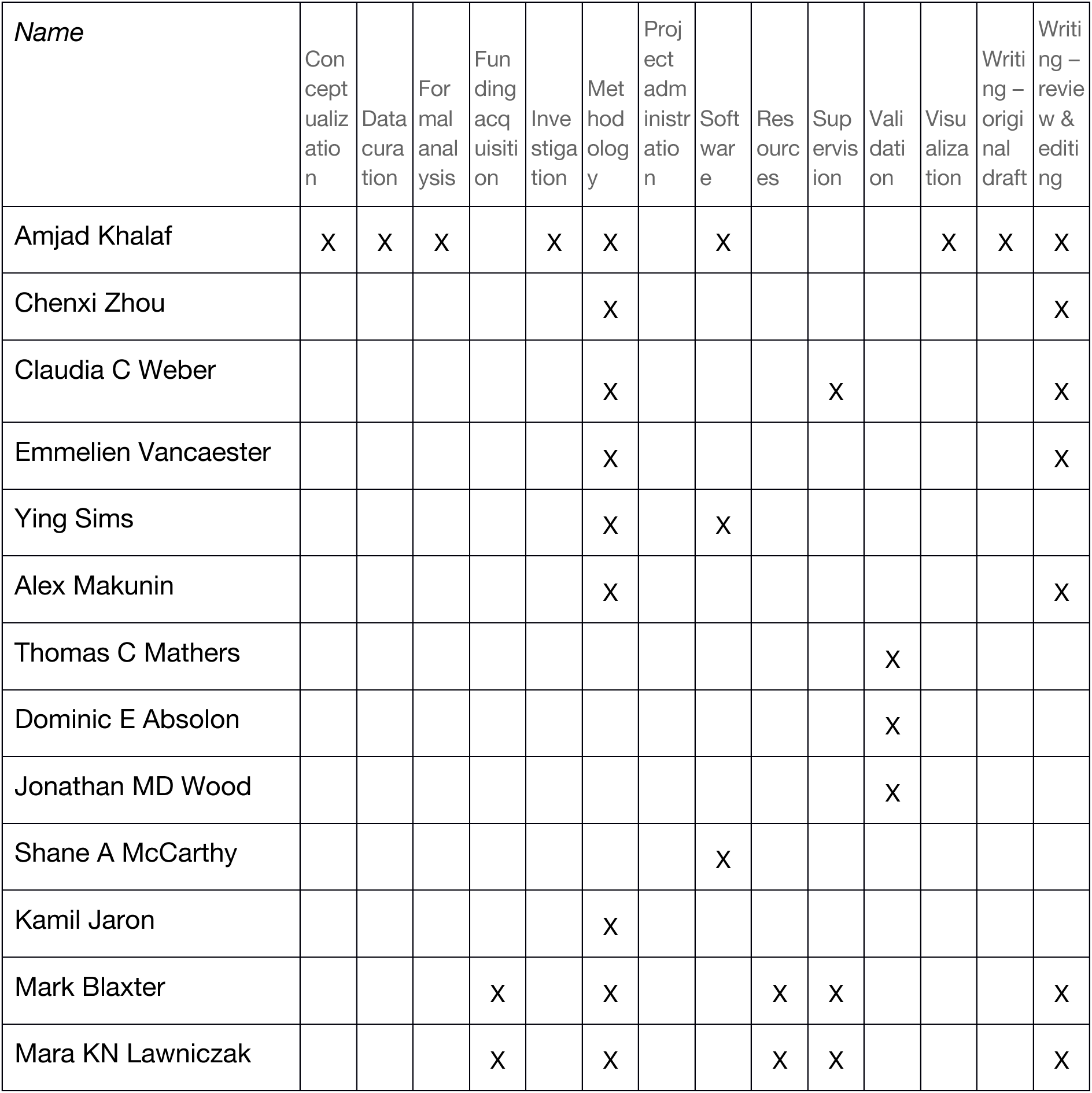

## References

1. Keeling P. Five questions about microsporidia. PLoS Pathog. 2009;5: e1000489. doi:10.1371/journal.ppat.1000489

2. Nageli C. uber die neue Krankheit der Seidenraupe und verwandte Organismen. [Abstract of report before 33. Versamml. Deutsch. Naturf. u. Aerzte. Bonn, 21 Sept.]. Bot Ztg. 1857;15: 760–761. Available: https://cir.nii.ac.jp/crid/1573105974684833920

3. Pasteur L. Etudes sur la maladie des vers à soie: 2.: Notes et documents. Gauthier-Villars; 1870. Available: https://play.google.com/store/books/details?id=y-1rmRQoAa4C

4. Bojko J, Reinke AW, Stentiford GD, Williams B, Rogers MSJ, Bass D. Microsporidia: a new taxonomic, evolutionary, and ecological synthesis. Trends Parasitol. 2022;38: 642– 659. doi:10.1016/j.pt.2022.05.007

5. Stentiford GD, Feist SW, Stone DM, Bateman KS, Dunn AM. Microsporidia: diverse, dynamic, and emergent pathogens in aquatic systems. Trends Parasitol. 2013;29: 567–578. doi:10.1016/j.pt.2013.08.005

6. Han B, Pan G, Weiss LM. Microsporidiosis in Humans. Clin Microbiol Rev. 2021;34: e0001020. doi:10.1128/CMR.00010-20

7. Becnel JJ, Weiss LM. Publication : USDA ARS. ars.usda.gov; 2014 [cited 12 Dec 2023]. Available: https://www.ars.usda.gov/research/publications/publication/?seqNo115=310041

8. Weber R, Bryan RT, Schwartz DA, Owen RL. Human microsporidial infections. Clin Microbiol Rev. 1994;7: 426–461. doi:10.1128/CMR.7.4.426

9. Bryan RT, Cali A, Owen RL, Spencer HC. Microsporidia: opportunistic pathogens in patients with AIDS. Prog Clin Parasitol. 1991;2: 1–26. Available: https://www.ncbi.nlm.nih.gov/pubmed/1893116

10. Matsubayashi H, Koike T, Mikata I, Takei H, Hagiwara S. A case of Encephalitozoon-like body infection in man. AMA Arch Pathol. 1959;67: 181–187. Available: https://www.ncbi.nlm.nih.gov/pubmed/13616827

11. Bojko J, Stentiford GD. Microsporidian Pathogens of Aquatic Animals. Experientia Suppl. 2022;114: 247–283. doi:10.1007/978-3-030-93306-7_10

12. Thomas SR, Elkinton JS. Pathogenicity and virulence. J Invertebr Pathol. 2004;85: 146–151. doi:10.1016/j.jip.2004.01.006

13. Anderson DL, Giacon H. Reduced pollen collection by honey bee (Hymenoptera: Apidae) colonies infected with Nosema apis and sacbrood virus. J Econ Entomol. 1992;85: 47–51. doi:10.1093/jee/85.1.47

14. Fries I, Ekbohm G, Villumstad E. Nosema Apis, Sampling Techniques and Honey Yield. J Apic Res. 1984;23: 102–105. doi:10.1080/00218839.1984.11100617

15. Vyas-Patel N. The Suppression of Plasmodium berghei in Anopheles coluzzii infected later with Vavraia culicis. bioRxiv. 2023. p. 2023.02.05.527158. doi:10.1101/2023.02.05.527158

16. Akorli J, Akorli EA, Tetteh SNA, Amlalo GK, Opoku M, Pwalia R, et al. Microsporidia MB is found predominantly associated with Anopheles gambiae s.s and Anopheles coluzzii in Ghana. Sci Rep. 2021;11: 18658. doi:10.1038/s41598-021-98268-2

17. Herren JK, Mbaisi L, Mararo E, Makhulu EE, Mobegi VA, Butungi H, et al. A microsporidian impairs Plasmodium falciparum transmission in Anopheles arabiensis mosquitoes. Nat Commun. 2020;11: 1–10. doi:10.1038/s41467-020-16121-y

18. Lee, Heitman, Ironside. Sex and the Microsporidia. Microsporidia: Pathogens of. 2014. Available: https://books.google.com/books?hl=en&lr=&id=k4kZBAAAQBAJ&oi=fnd&pg=PA231&dq=microsporidia+ploidy&ots=oqF0tRPXyc&sig=Ua18f4MX3vgjvbaipK2DYB1v3Kc

19. Amigó JM, Gracia MP, Salvadó H, Vivarés CP. Pulsed Field Gel Electrophoresis of Three Microsporidian Parasites of Fish. Acta Protozool. 2002;41: 11–16. Available: https://www.airitilibrary.com/Publication/alDetailedMesh?docid=00651583-200203-201012290004-201012290004-11-16

20. Bernander R, Palm JE, Svärd SG. Genome ploidy in different stages of the Giardia lamblia life cycle. Cell Microbiol. 2001;3: 55–62. doi:10.1046/j.1462-5822.2001.00094.x

21. Vávra J. Development of the Microsporidia. In: Bulla LA, Cheng TC, editors. Biology of the Microsporidia. Boston, MA: Springer US; 1976. pp. 87–109. doi:10.1007/978-1-4684-3114-8_3

22. Youssef NN, Hammond DM. The fine structure of the developmental stages of the microsporidian Nosema apis Zander. Tissue Cell. 1971;3: 283–294. doi:10.1016/s0040-8166(71)80023-x

23. Cali A. Morphogenesis in the genus Nosema. ProcIVth IntColloqInsect Pathol. 1971;4: 104–112. Available: https://ci.nii.ac.jp/naid/10004755181/

24. Sprague V, Vernick SH. The ultrastructure of Encephalitozoon cuniculi (Microsporida, Nosematidae) and its taxonomic significance. J Protozool. 1971;18: 560–569. doi:10.1111/j.1550-7408.1971.tb03376.x

25. Miquel J, Kacem H, Baz-González E, Foronda P, Marchand B. Ultrastructural and molecular study of the microsporidian Toguebayea baccigeri n. gen., n. sp., a hyperparasite of the digenean trematode Bacciger israelensis (Faustulidae), a parasite of Boops boops (Teleostei, Sparidae). Parasite. 2022;29: 2. doi:10.1051/parasite/2022007

26. Pretto T, Montesi F, Ghia D, Berton V, Abbadi M, Gastaldelli M, et al. Ultrastructural and molecular characterization of Vairimorpha austropotamobii sp. nov. (Microsporidia: Burenellidae) and Thelohania contejeani (Microsporidia: Thelohaniidae), two parasites of the white-clawed crayfish, Austropotamobius pallipes complex (Decapoda: Astacidae). J Invertebr Pathol. 2018;151: 59–75. doi:10.1016/j.jip.2017.11.002

27. Maurand J, Vey A. Etudes histopathologique et ultrastructurale de Thelohania contejeani (Microsporida, Nosematidae) parasite de l’Ecrevisse Austropotamobius pallipes Lereboullet. Ann Parasitol Hum Comp. 1973;48: 411–421. doi:10.1051/parasite/1973483411

28. Weidner E. Ultrastructural study of microsporidian development. Zeitschrift für Zellforschung und Mikroskopische Anatomie. 1970;105: 33–54. doi:10.1007/BF00340563

29. Sokolova YY, Fuxa JR. Biology and life-cycle of the microsporidium Kneallhazia solenopsae Knell Allan Hazard 1977 gen. n., comb. n., from the fire ant Solenopsis invicta. Parasitology. 2008;135: 903–929. doi:10.1017/S003118200800440X

30. Becnel JJ. Horizontal transmission and subsequent development of Amblyospora californica (Microsporida: Amblyosporidae) in the intermediate and definitive hosts. Dis Aquat Organ. 1992;13: 17–28. Available: https://www.int-res.com/articles/dao/13/d013p017.pdf

31. Becnel JJ, Sprague V, Fukuda T, Hazard EI. Development of Edhazardia aedis (Kudo, 1930) n. g., n. comb. (Microsporida: Amblyosporidae) in the mosquito Aedes aegypti (L.) (Diptera: Culicidae). J Protozool. 1989;36: 119–130. doi:10.1111/j.1550-7408.1989.tb01057.x

32. Canning EU. Nuclear division and chromosome cycle in microsporidia. Biosystems. 1988;21: 333–340. doi:10.1016/0303-2647(88)90030-5

33. Becnel JJ, Hazard EI, Fukuda T, Sprague V. Life cycle ofCulicospora magna(Kudo, 1920) (microsporida: Culicosporidae) inCulex restuansTheobald with special reference to Sexuality1. J Protozool. 1987;34: 313–322. doi:10.1111/j.1550-7408.1987.tb03182.x

34. Hazard EI, Fukuda T, Becnel JJ. Gametogenesis and plasmogamy in certain species of Microspora. J Invertebr Pathol. 1985;46: 63–69. doi:10.1016/0022-2011(85)90130-2

35. Hazard EI, Brookbank JW. Karyogamy and meiosis in an Amblyospora sp. (Microspora) in the mosquito Culex salinarius. J Invertebr Pathol. 1984;44: 3–11. doi:10.1016/0022-2011(84)90039-9

36. Hazard EI, Andreadis TG, Joslyn DJ, Ellis EA. Meiosis and Its Implications in the Life Cycles of Amblyospora and Parathelohania (Microspora). J Parasitol. 1979;65: 117–122. doi:10.2307/3280215

37. Vivares CP, Sprague V. The fine structure of Ameson pulvis (Microspora, Microsporida) and its implications regarding classification and chromosome cycle. J Invertebr Pathol. 1979;33: 40–52. doi:10.1016/0022-2011(79)90128-9

38. Loubès C. [Meiosis in Microsporidia: effects on biological cycles]. J Protozool. 1979;26: 200–208. doi:10.1111/j.1550-7408.1979.tb02761.x

39. Loubès C, Maurand J, Rousset-Galangau V. [Presence of synaptonematic complexes in the biological cycle of Gurleya chironomi Loubes and Maurand, 1975: an argument in favor of sexuality in microsporidia]. C R Acad Hebd Seances Acad Sci D. 1976;282: 1025–1027. Available: https://www.ncbi.nlm.nih.gov/pubmed/821628

40. Desportes I. ULTRASTRUCTURE DE STEMPELLIA MUTABILIS LEGER ET HESSE, MICROSPORIDIE PARASITE DE L’EPHEMERE EPHEMERA VULGATA L. 1976 [cited 29 Dec 2022]. Available: https://pascal-francis.inist.fr/vibad/index.php?action=getRecordDetail&idt=PASCAL7750051096

41. Cali A, Becnel JJ, Takvorian PM. Microsporidia. In: Archibald JM, Simpson AGB, Slamovits CH, editors. Handbook of the Protists. Cham: Springer International Publishing; 2017. pp. 1559–1618. doi:10.1007/978-3-319-28149-0_27

42. Sokolova YY, Dolgikh VV, Morzhina EV, Nassonova ES, Issi IV, Terry RS, et al. Establishment of the new genus Paranosema based on the ultrastructure and molecular phylogeny of the type species Paranosema grylli Gen. Nov., Comb. Nov. (Sokolova, Selezniov, Dolgikh, Issi 1994), from the cricket Gryllus bimaculatus Deg. J Invertebr Pathol. 2003;84: 159–172. doi:10.1016/j.jip.2003.10.004

43. Nassonova ES, Smirnov AV. Synaptonemal complexes as evidence for meiosis in the life cycle of the monomorphic diplokaryotic microsporidium Paranosema grylli. Eur J Protistol. 2005;41: 175–181. doi:10.1016/j.ejop.2005.02.001

44. Angst P, Ebert D, Fields PD. Demographic history shapes genomic variation in an intracellular parasite with a wide geographical distribution. Mol Ecol. 2022;31: 2528– 2544. doi:10.1111/mec.16419

45. Cuomo CA, Desjardins CA, Bakowski MA, Goldberg J, Ma AT, Becnel JJ, et al. Microsporidian genome analysis reveals evolutionary strategies for obligate intracellular growth. Genome Res. 2012;22: 2478–2488. doi:10.1101/gr.142802.112

46. Khalaf A, Lawniczak MKN, Blaxter ML, Jaron KS. Polyploidy is widespread in Microsporidia. Microbiol Spectr. 2024;12: e0366923. doi:10.1128/spectrum.03669-23

47. Stratton CE, Bolds SA, Reisinger LS, Behringer DC, Khalaf A, Bojko J. Microsporidia and invertebrate hosts: genome-informed taxonomy surrounding a new lineage of crayfish-infecting Nosema spp. (Nosematida). Fungal Divers. 2024;128: 167–190. doi:10.1007/s13225-024-00543-w

48. Pelin A, Selman M, Aris-Brosou S, Farinelli L, Corradi N. Genome analyses suggest the presence of polyploidy and recent human-driven expansions in eight global populations of the honeybee pathogen Nosema ceranae. Environ Microbiol. 2015;17: 4443–4458. doi:10.1111/1462-2920.12883

49. Sokolova YY, Overstreet RM. A new microsporidium, Apotaspora heleios n. g., n. sp., from the Riverine grass shrimp Palaemonetes paludosus (Decapoda: Caridea: Palaemonidae). J Invertebr Pathol. 2018;157: 125–135. doi:10.1016/j.jip.2018.05.007

50. Otto SP. The evolutionary consequences of polyploidy. Cell. 2007;131: 452–462. doi:10.1016/j.cell.2007.10.022

51. Spoelhof JP, Soltis PS, Soltis DE. Pure polyploidy: Closing the gaps in autopolyploid research: Pure polyploidy. J Syst Evol. 2017;55: 340–352. doi:10.1111/jse.12253

52. Redmond AK, Casey D, Gundappa MK, Macqueen DJ, McLysaght A. Independent rediploidization masks shared whole genome duplication in the sturgeon-paddlefish ancestor. Nat Commun. 2023;14: 2879. doi:10.1038/s41467-023-38714-z

53. Li Z, McKibben MTW, Finch GS, Blischak PD, Sutherland BL, Barker MS. Patterns and processes of diploidization in land plants. Annu Rev Plant Biol. 2021;72: 387–410. doi:10.1146/annurev-arplant-050718-100344

54. Du K, Stöck M, Kneitz S, Klopp C, Woltering JM, Adolfi MC, et al. The sterlet sturgeon genome sequence and the mechanisms of segmental rediploidization. Nat Ecol Evol. 2020;4: 841–852. doi:10.1038/s41559-020-1166-x

55. Mandáková T, Lysak MA. Post-polyploid diploidization and diversification through dysploid changes. Curr Opin Plant Biol. 2018;42: 55–65. doi:10.1016/j.pbi.2018.03.001

56. Robertson FM, Gundappa MK, Grammes F, Hvidsten TR, Redmond AK, Lien S, et al. Lineage-specific rediploidization is a mechanism to explain time-lags between genome duplication and evolutionary diversification. Genome Biol. 2017;18: 111. doi:10.1186/s13059-017-1241-z

57. Conant GC, Birchler JA, Pires JC. Dosage, duplication, and diploidization: clarifying the interplay of multiple models for duplicate gene evolution over time. Curr Opin Plant Biol. 2014;19: 91–98. doi:10.1016/j.pbi.2014.05.008

58. Hokamp K, McLysaght A, Wolfe KH. The 2R hypothesis and the human genome sequence. J Struct Funct Genomics. 2003;3: 95–110. Available: https://pubmed.ncbi.nlm.nih.gov/12836689/

59. Furlong RF, Holland PWH. Were vertebrates octoploid? Philos Trans R Soc Lond B Biol Sci. 2002;357: 531–544. doi:10.1098/rstb.2001.1035

60. Wolfe KH. Yesterday’s polyploids and the mystery of diploidization. Nat Rev Genet. 2001;2: 333–341. doi:10.1038/35072009

61. Khalaf A, Francis O, Blaxter ML. Genome evolution in intracellular parasites: Microsporidia and Apicomplexa. J Eukaryot Microbiol. 2024; e13033. doi:10.1111/jeu.13033

62. The Darwin Tree of Life Project Consortium, Blaxter M, Mieszkowska N, Palma FD, Holland P, Durbin R, et al. Sequence locally, think globally: The Darwin Tree of Life Project. Proceedings of the National Academy of Sciences. 2022;119: e2115642118. doi:10.1073/pnas.2115642118

63. Weber CC, Paulini M, Wellcome Sanger Institute Tree of Life Management, Samples and Laboratory team, Wellcome Sanger Institute Tree of Life Core Informatics team, Blaxter ML. *Kudoa*genomes from contaminated hosts reveal extensive gene order conservation and rapid sequence evolution. bioRxiv. 2024. p. 2024.11.01.621499. doi:10.1101/2024.11.01.621499

64. Vancaester E, Blaxter M. Phylogenomic analysis of Wolbachia genomes from the Darwin Tree of Life biodiversity genomics project. PLoS Biol. 2023;21: e3001972. doi:10.1371/journal.pbio.3001972

65. Weber CC. Disentangling cobionts and contamination in long-read genomic data using sequence composition. G3 (Bethesda). 2024;14: jkae187. doi:10.1093/g3journal/jkae187

66. Vancaester E, Blaxter ML. MarkerScan: Separation and assembly of cobionts sequenced alongside target species in biodiversity genomics projects. Wellcome Open Res. 2024;9: 33. doi:10.12688/wellcomeopenres.20730.1

67. Mautner SI, Cook KA, Forbes MR, McCurdy DG, Dunn AM. Evidence for sex ratio distortion by a new microsporidian parasite of a Corophiid amphipod. Parasitology. 2007;134: 1567–1573. doi:10.1017/S0031182007003034

68. Desjardins CA, Sanscrainte ND, Goldberg JM, Heiman D, Young S, Zeng Q, et al. Contrasting host-pathogen interactions and genome evolution in two generalist and specialist microsporidian pathogens of mosquitoes. Nat Commun. 2015;6: 7121. doi:10.1038/ncomms8121

69. Zhang C, Rabiee M, Sayyari E, Mirarab S. ASTRAL-III: polynomial time species tree reconstruction from partially resolved gene trees. BMC Bioinformatics. 2018;19: 153. doi:10.1186/s12859-018-2129-y

70. Simão FA, Waterhouse RM, Ioannidis P, Kriventseva EV, Zdobnov EM. BUSCO: assessing genome assembly and annotation completeness with single-copy orthologs. Bioinformatics. 2015;31: 3210–3212. doi:10.1093/bioinformatics/btv351

71. Minh BQ, Schmidt HA, Chernomor O, Schrempf D, Woodhams MD, von Haeseler A, et al. IQ-TREE 2: New Models and Efficient Methods for Phylogenetic Inference in the Genomic Era. Mol Biol Evol. 2020;37: 1530–1534. doi:10.1093/molbev/msaa015

72. Seemann T. Prokka: rapid prokaryotic genome annotation. Bioinformatics. 2014;30: 2068–2069. doi:10.1093/bioinformatics/btu153

73. Flynn JM, Hubley R, Goubert C, Rosen J, Clark AG, Feschotte C, et al. RepeatModeler2 for automated genomic discovery of transposable element families. Proceedings of the National Academy of Sciences. 2020;117: 9451–9457. doi:10.1073/pnas.1921046117

74. Smit AFA, Hubley R, Green P. RepeatMasker Open-4.0.2013-2015. 2013. Available: <http://www.repeatmasker.org>

75. Pagel M. Detecting correlated evolution on phylogenies: a general method for the comparative analysis of discrete characters. Proceedings of the Royal Society of London Series B: Biological Sciences. 1994;255: 37–45. doi:10.1098/rspb.1994.0006

76. Cormier A, Wattier R, Giraud I, Teixeira M, Grandjean F, Rigaud T, et al. Draft Genome Sequences of Thelohania contejeani and Cucumispora dikerogammari, Pathogenic Microsporidia of Freshwater Crustaceans. Microbiol Resour Announc. 2021;10. doi:10.1128/MRA.01346-20

77. Wadi L, Reinke AW. Evolution of microsporidia: An extremely successful group of eukaryotic intracellular parasites. PLoS Pathog. 2020;16: e1008276. doi:10.1371/journal.ppat.1008276

78. Bartolomé C, Higes M, Hernández RM, Chen YP, Evans JD, Huang Q. The recent revision of the genera Nosema and Vairimorpha (Microsporidia: Nosematidae) was flawed and misleads the bee scientific community. J Invertebr Pathol. 2024;206: 108146. doi:10.1016/j.jip.2024.108146

79. Tokarev YS, Huang W-F, Solter LF, Malysh JM, Becnel JJ, Vossbrinck CR. A formal redefinition of the genera Nosema and Vairimorpha (Microsporidia: Nosematidae) and reassignment of species based on molecular phylogenetics. J Invertebr Pathol. 2020;169: 107279. doi:10.1016/j.jip.2019.107279

80. Birky CW Jr. Heterozygosity, heteromorphy, and phylogenetic trees in asexual eukaryotes. Genetics. 1996;144: 427–437. doi:10.1093/genetics/144.1.427

81. Shimodaira H. An approximately unbiased test of phylogenetic tree selection. Syst Biol. 2002;51: 492–508. doi:10.1080/10635150290069913

82. Barton AB, Pekosz MR, Kurvathi RS, Kaback DB. Meiotic recombination at the ends of chromosomes in Saccharomyces cerevisiae. Genetics. 2008;179: 1221–1235. doi:10.1534/genetics.107.083493

83. Jensen-Seaman MI, Furey TS, Payseur BA, Lu Y, Roskin KM, Chen C-F, et al. Comparative recombination rates in the rat, mouse, and human genomes. Genome Res. 2004;14: 528–538. doi:10.1101/gr.1970304

84. Li H. Minimap2: pairwise alignment for nucleotide sequences. Bioinformatics. 2018;34: 3094–3100. doi:10.1093/bioinformatics/bty191

85. PretextView: OpenGL Powered Pretext Contact Map Viewer. Github; Available: https://github.com/sanger-tol/PretextView

86. Chen H, Zwaenepoel A, Van de Peer Y. Wgd v2: A suite of tools to uncover and date ancient polyploidy and whole-genome duplication. Bioinformatics. 2024;40: btae272. doi:10.1093/bioinformatics/btae272

87. Maere S, De Bodt S, Raes J, Casneuf T, Van Montagu M, Kuiper M, et al. Modeling gene and genome duplications in eukaryotes. Proc Natl Acad Sci U S A. 2005;102: 5454–5459. doi:10.1073/pnas.0501102102

88. Van de Peer Y. Computational approaches to unveiling ancient genome duplications. Nat Rev Genet. 2004;5: 752–763. doi:10.1038/nrg1449

89. Blanc G, Wolfe KH. Widespread paleopolyploidy in model plant species inferred from age distributions of duplicate genes. Plant Cell. 2004;16: 1667–1678. doi:10.1105/tpc.021345

90. Lynch M, Conery JS. The evolutionary demography of duplicate genes. Genome Evolution. Dordrecht: Springer Netherlands; 2003. pp. 35–44. doi:10.1007/978-94-010-0263-9_4

91. Ranallo-Benavidez TR, Jaron KS, Schatz MC. GenomeScope 2.0 and Smudgeplot for reference-free profiling of polyploid genomes. Nat Commun. 2020;11: 1–10. doi:10.1038/s41467-020-14998-3

92. Marçais G, Kingsford C. A fast, lock-free approach for efficient parallel counting of occurrences of k-mers. Bioinformatics. 2011;27: 764–770. doi:10.1093/bioinformatics/btr011

93. Mackintosh A, de la Rosa PMG, Martin SH, Lohse K, Laetsch DR. Inferring inter-chromosomal rearrangements and ancestral linkage groups from synteny. bioRxiv. 2023. p. 2023.09.17.558111. doi:10.1101/2023.09.17.558111

94. Pedregosa F, Varoquaux G, Gramfort A, Michel V, Thirion B, Grisel O, et al. Scikit-learn: Machine Learning in Python. J Mach Learn Res. 2011;12: 2825–2830. doi:10.5555/1953048.2078195

95. Maaten L, Hinton GE. Visualizing Data using t-SNE. Journal of Machine Learning Research. 2008;9: 2579–2605. Available: https://www.jmlr.org/papers/volume9/vandermaaten08a/vandermaaten08a.pdf

96. Schultz DT, Haddock SHD, Bredeson JV, Green RE, Simakov O, Rokhsar DS. Ancient gene linkages support ctenophores as sister to other animals. Nature. 2023;618: 110–117. doi:10.1038/s41586-023-05936-6

97. Konstantinidis KT, Tiedje JM. Genomic insights that advance the species definition for prokaryotes. Proc Natl Acad Sci U S A. 2005;102: 2567–2572. doi:10.1073/pnas.0409727102

98. Hart R, Moran NA, Ochman H. Genomic divergence across the tree of life. Proc Natl Acad Sci U S A. 2025;122: e2319389122. doi:10.1073/pnas.2319389122

99. Ohno S. Evolution by gene duplication. 1970. Available: https://books.google.com/books?hl=en&lr=&id=5SjqCAAAQBAJ&oi=fnd&pg=PA1&ots=MoU4vLG0Af&sig=7ZznL61U389lqqWhblOVWnIAK-k

100. Szadkowski E, Eber F, Huteau V, Lodé M, Huneau C, Belcram H, et al. The first meiosis of resynthesized Brassica napus, a genome blender. New Phytol. 2010;186: 102–112. doi:10.1111/j.1469-8137.2010.03182.x

101. Udall JA, Quijada PA, Osborn TC. Detection of chromosomal rearrangements derived from homologous recombination in four mapping populations of Brassica napus L. Genetics. 2005;169: 967–979. doi:10.1534/genetics.104.033209

102. Schranz ME, Osborn TC. De novo variation in life-history traits and responses to growth conditions of resynthesized polyploid Brassica napus (Brassicaceae). Am J Bot. 2004;91: 174–183. doi:10.3732/ajb.91.2.174

103. Osborn TC, Butrulle DV, Sharpe AG, Pickering KJ, Parkin IAP, Parker JS, et al. Detection and effects of a homeologous reciprocal transposition in Brassica napus. Genetics. 2003;165: 1569–1577. doi:10.1093/genetics/165.3.1569

104. Song K, Lu P, Tang K, Osborn TC. Rapid genome change in synthetic polyploids of Brassica and its implications for polyploid evolution. Proc Natl Acad Sci U S A. 1995;92: 7719–7723. doi:10.1073/pnas.92.17.7719

105. Weiss H, Maluszynska J. Chromosomal rearrangement in autotetraploid plants of Arabidopsis thaliana. Hereditas. 2000;133: 255–261. doi:10.1111/j.1601-5223.2000.00255.x

106. Nakao Y, Kanamori T, Itoh T, Kodama Y, Rainieri S, Nakamura N, et al. Genome sequence of the lager brewing yeast, an interspecies hybrid. DNA Res. 2009;16: 115–129. doi:10.1093/dnares/dsp003

107. Usher J, Bond U. Recombination between homoeologous chromosomes of lager yeasts leads to loss of function of the hybrid GPH1 gene. Appl Environ Microbiol. 2009;75: 4573–4579. doi:10.1128/AEM.00351-09

108. Dunn B, Sherlock G. Reconstruction of the genome origins and evolution of the hybrid lager yeast Saccharomyces pastorianus. Genome Res. 2008;18: 1610–1623. doi:10.1101/gr.076075.108

109. Shin H, Park JE, Park HR, Choi WL, Yu SH, Koh W, et al. Admixture of divergent genomes facilitates hybridization across species in the family Brassicaceae. New Phytol. 2022;235: 743–758. doi:10.1111/nph.18155

110. McClintock B. The significance of responses of the genome to challenge. Science. 1984;226: 792–801. doi:10.1126/science.15739260

111. Storchová Z, Breneman A, Cande J, Dunn J, Burbank K, O’Toole E, et al. Genome-wide genetic analysis of polyploidy in yeast. Nature. 2006;443: 541–547. doi:10.1038/nature05178

112. Andreadis TG. Amblyospora connecticus sp. nov. (Microsporida: Amblyosporidae): Horizontal transmission studies in the mosquito Aedes cantator and formal description. J Invertebr Pathol. 1988;52: 90–101. doi:10.1016/0022-2011(88)90107-3

113. Murareanu BM, Sukhdeo R, Qu R, Jiang J, Reinke AW. Generation of a Microsporidia Species Attribute Database and Analysis of the Extensive Ecological and Phenotypic Diversity of Microsporidia. MBio. 2021;12: e0149021. doi:10.1128/mBio.01490-21

114. Cali A, Takvorian PM. Developmental morphology and life cycles of the microsporidia. Microsporidia: Pathogens of opportunity. 2014. Available: https://onlinelibrary.wiley.com/doi/abs/10.1002/9781118395264.ch2

115. Baker MD, Vossbrinck CR, Didier ES, Maddox JV, Shadduck JA. Small subunit ribosomal DNA phylogeny of various microsporidia with emphasis on AIDS related forms. J Eukaryot Microbiol. 1995;42: 564–570. doi:10.1111/j.1550-7408.1995.tb05906.x

116. Zhu X, Wittner M, Tanowitz HB, Kotler D, Cali A, Weiss LM. Small subunit rRNA sequence of Enterocytozoon bieneusi and its potential diagnostic role with use of the polymerase chain reaction. J Infect Dis. 1993;168: 1570–1575. doi:10.1093/infdis/168.6.1570

117. Cheng H, Concepcion GT, Feng X, Zhang H, Li H. Haplotype-resolved de novo assembly using phased assembly graphs with hifiasm. Nat Methods. 2021;18: 170–175. doi:10.1038/s41592-020-01056-5

118. Challis R, Richards E, Rajan J, Cochrane G, Blaxter M. BlobToolKit – Interactive Quality Assessment of Genome Assemblies. G3 Genes|Genomes|Genetics. 2020;10: 1361–1374. doi:10.1534/g3.119.400908

119. Li H, Handsaker B, Wysoker A, Fennell T, Ruan J, Homer N, et al. The Sequence Alignment/Map format and SAMtools. Bioinformatics. 2009;25: 2078–2079. doi:10.1093/bioinformatics/btp352

120. Vasimuddin M, Misra S, Li H, Aluru S. Efficient architecture-aware acceleration of BWA-MEM for multicore systems. 2019 IEEE International Parallel and Distributed Processing Symposium (IPDPS). IEEE; 2019. pp. 314–324. doi:10.1109/ipdps.2019.00041

121. Dudchenko O, Shamim MS, Batra SS, Durand NC, Musial NT, Mostofa R, et al. The Juicebox Assembly Tools module facilitates de novo assembly of mammalian genomes with chromosome-length scaffolds for under $1000. bioRxiv. bioRxiv; 2018. p. 254797. doi:10.1101/254797

122. Quinlan AR, Hall IM. BEDTools: a flexible suite of utilities for comparing genomic features. Bioinformatics. 2010;26: 841–842. doi:10.1093/bioinformatics/btq033

123. PretextMap: Paired REad TEXTure Mapper. Converts SAM formatted read pairs into genome contact maps. Github; Available: https://github.com/sanger-tol/PretextMap

124. Guan D, McCarthy SA, Wood J, Howe K, Wang Y, Durbin R. Identifying and removing haplotypic duplication in primary genome assemblies. Bioinformatics. 2020;36: 2896–2898. doi:10.1093/bioinformatics/btaa025

125. Katoh K, Misawa K, Kuma K, Miyata T. MAFFT: a novel method for rapid multiple sequence alignment based on fast Fourier transform. Nucleic Acids Res. 2002;30: 3059–3066. doi:10.1093/nar/gkf436

126. Tournayre J, Polonais V, Wawrzyniak I, Akossi RF, Parisot N, Lerat E, et al. MicroAnnot: A dedicated workflow for accurate microsporidian genome annotation. Int J Mol Sci. 2024;25. doi:10.3390/ijms25020880

127. Whelan TA, Lee NT, Lee RCH, Fast NM. Microsporidian introns retained against a background of genome reduction: Characterization of an unusual set of introns. Genome Biol Evol. 2019;11: 263–269. doi:10.1093/gbe/evy260

