## Supporting Information Guide for "Forty New Genomes Shed Light on Sexual Reproduction and the Origin of Tetraploidy in Microsporidia"

This PDF file includes:

- The list and legend descriptions of all the files and materials contained in this Supporting Information dataset.
- Supporting Information Figures 1 to 9
- Supporting Information Tables 4 and 5

Find Supporting Information in full on Zenodo https://doi.org/10.5281/zenodo.15364389.

Author information


| *Name* | *ORCID* | *email* |
|---|---|---|
| Amjad Khalaf | 0000-0003-1297-1181 | |
| Chenxi Zhou | 0000-0002-1735-2630 | |
| Claudia C Weber | 0000-0002-5910-8898 | |
| Emmelien Vancaester | 0000-0002-9177-8808 | |
| Ying Sims | 0000-0003-4765-4872 | |
| Alex Makunin | 0000-0002-9555-5097 | |
| Thomas C Mathers | 0000-0002-8637-3515 | |
| Dominic E Absolon | 0009-0004-5709-2249 | |
| Jonathan MD Wood | 0000-0002-7545-2162 | |
| Shane A McCarthy | 0000-0002-2715-4187 | |
| Kamil Jaron | 0000-0003-1470-5450 | |
| Mark Blaxter | 0000-0003-2861-949X | |
| Mara KN Lawniczak | 0000-0002-3006-2080 | |

#### Author contributions

| *Name* | Conceptualization | Data curation | Formal analysis | Funding acquisition | Investigation | Methodology | Project administration | Software | Resources | Supervision | Validation | Visualization | Writing – original draft | Writing – review & editing |
|---|---|---|---|---|---|---|---|---|---|---|---|---|---|---|
| Amjad Khalaf | X | X | X | | X | X | | X | | | | X | X | X |
| Chenxi Zhou | | | | | | X | | | | | | | | X |
| Claudia C Weber | | | | | | X | | | | X | | | | X |
| Emmelien Vancaester | | | | | | X | | | | | | | | X |
| Ying Sims | | | | | | X | | X | | | | | | |
| Alex Makunin | | | | | | X | | | | | | | | X |
| Thomas C Mathers | | | | | | | | | | | X | | | |
| Dominic E Absolon | | | | | | | | | | | X | | | |
| Jonathan MD Wood | | | | | | | | | | | X | | | |
| Shane A McCarthy | | | | | | | | X | | | | | | |
| Kamil Jaron | | | | | | X | | | | | | | | |
| Mark Blaxter | | | | X | | X | | | X | X | | | | X |
| Mara KN Lawniczak | | | | X | | X | | | X | X | | | | X |

### Supporting Information Section 1

Table S1: Microsporidian genome assemblies.

Full list of recovered microsporidian genome assemblies and their associated meta-data. Genome accessions will be added as genomes are released through ENA.

**Zenodo: Supporting Information Table S1**

File Collection S1: Microsporidian genome assemblies fasta files.

Fasta files of recovered microsporidian genome assemblies. The primary assemblies listed in Table S1 are given by {Host ToLID}.µ.fasta, whereas purged haplotypic duplication sequences are given by {Host ToLID}.µ.alt.fasta where applicable. In the case of iuLoeVari1.µ, the primary assembly is the best haploid representative genome assembly possible, containing sequences across all four compartments. The diploid genome assemblies of iuLoeVari1.µ's AB and CD compartments are given by iuLoeVari1.µ.AB.fasta and iuLoeVari1.µ.CD.fasta respectively.

**Zenodo: Supporting Information File Collection S1**

Table S2: Filtering parameters used in generating genome assemblies.

Parameters used for filtering microsporidian contigs from their respective (meta-)genomic assemblies in filtering steps 1 (BlobToolKit [1]) and 2 (BubblePlot, Github: https://github.com/Amjad-Khalaf/BubblePlot). See Materials and Methods for details.

**Zenodo: Supporting Information Table S2**

#### File Collection S2: Statistics of intermediate steps for each microsporidian genome assembly.

Scaffold/contig and read statistics for intermediate steps produced in the generation of each microsporidian genome assembly.

**Zenodo: Supporting Information File Collection S2**

#### File Collection S3: K-mer analysis plots for the microsporidian genome assemblies.

MerqurkyFK plots (Github: https://github.com/thegenemyers/MERQURY.FK) for final microsporidian genome assemblies generated in this study.

**Zenodo: Supporting Information File Collection S3**

#### File Collection S4: K-mer histogram plots for the reads used to produce the microsporidian genome assemblies.

GenomeScope2 [2] plots of the reads used to produce the microsporidian genome assemblies presented in this study. Jellyfish was used to generate the k-mer spectrum for each read set (k = 21, version 2.2.10) [3].

**Zenodo: Supporting Information File Collection S4**

#### File Collection S5: Smudgeplot ploidy estimation.

Smudgeplot [2] plots of the reads used to produce the microsporidian genome assemblies for which ploidy could be estimated using GenomeScope2 [2] (Supplementary Information Fig. S41-S81).

**Zenodo: Supporting Information File Collection S5**

File Collection S6: K-mer plots used to inform genome assembly purging.

Purge_dups histogram plots used to inform genome assembly purging, with cutoffs used clearly indicated.

**Zenodo: Supporting Information File Collection S6**

File Collection S7: Self alignment plots for microsporidian genome assemblies.

Self-alignment dot plots for the microsporidian genome assemblies generated in this study. Each genome was aligned to itself using FASTGA (Github: https://github.com/thegenemyers/FASTGA). The plots were generated using HyraxDotPlot (v2.0) (Github: https://github.com/Amjad-Khalaf/HyraxDotPlot).

**Zenodo: Supporting Information File Collection S7**

File Collection S8: Hi-C contact heatmaps for scaffolded microsporidian genome assemblies.

Hi-C contact heatmaps for scaffolded microsporidian genome assemblies visualised using PretextView [4].

**Zenodo: Supporting Information File Collection S8**

File Collection S9: Oxford Dot Plots for tetraploid microsporidian genome assemblies.

Oxford dot plot for tetraploid genome assemblies displaying BUSCO genes. Gene pairs which are less divergent than the same species threshold are in sky blue, while gene pairs which are more divergent than the same species threshold are in red.

**Zenodo: Supporting Information File Collection S9**

File Collection S10: Annotation results for microsporidian genome assemblies.

Results for gene annotation (with Prokka) and repeat annotation (with RepeatModeler and RepeatMasker) on all genomes [5–7].

**Zenodo: Supporting Information File Collection S10**

Table S3: Accession numbers for publicly available genomes used in this study.

On the 1st of January 2025, we downloaded all microsporidian genome assemblies available in the NCBI Genome database. This retrieved 106 genome assemblies.

**Zenodo: Supporting Information Table S3**

### Supporting Information Section 2

Table S4: Trait-phylogeny regression.

Transformations representing the fit with the tree’s topology (λ), branch-lengths (κ) and root-tip distance (δ) [8] and the number of coding sequences, transposable element loads, and genome spans.

| | Tree topology (λ) | Branch-lengths (κ) | Root-tip distance (δ) |
|---|---|---|---|
| Genome span | 0.000 | 0.000 | 0.000 |
| Number of coding sequences | 0.000 | 0.000 | 0.000 |
| Retroelement load | 0.000 | 0.020 | 1.14 |
| DNA transposon load | 0.000 | 0.000 | 3.000 |
| Helitron load | 0.000 | 0.593 | 1.143 |

Table S5: Trait correlation.

Correlations between the number of coding sequences, transposable element loads, and genome spans.

| | Genome span | Number of coding sequences | Retroelement load | DNA transposon load | Helitron load |
|---|---|---|---|---|---|
| Genome span | 1.000 | 0.946 | 0.507 | 0.492 | 0.173 |
| Number of coding sequences | 0.946 | 1.000 | 0.462 | 0.556 | 0.196 |
| Retroelement | 0.507 | 0.462 | 1.000 | 0.413 | 0.104 |

| | | | | | |
|---|---|---|---|---|---|
| load | | | | | |
| DNA transposon load | 0.492 | 0.556 | 0.413 | 1.000 | 0.214 |
| Helitron load | 0.173 | 0.196 | 0.104 | 0.214 | 1.000 |

### Supporting Information Section 3

Table S6: Branch length distances for species delineation.

Pairwise branch length distances which include one of our genomes, and can be classified to a species or a genus. The conservative branch length threshold range was defined using the shortest observed branch lengths between known same-species genomes for the lower bound (0) and the smallest distance between *H*. *tvaerminnensis* and *H*. *magnivora* genomes for the upper bound (0.012). The relaxed threshold uses the full range of observed branch lengths among known same-species genomes (excluding the *H*. *tvaerminnensis* – *H*. *magnivora* cutoff).

**Zenodo: Supporting Information Table S6**

Fig. S1: Comparison of whole-genome phylogeny species delineation thresholds and individual gene phylogeny branch length distribution species delineation thresholds.

The approach we presented in Fig. 3 relies on branch lengths derived from the whole-genome phylogeny in Fig. 2 (i.e. a concatenated supermatrix of genes). We re-estimated same-species branch length thresholds for each gene. For each gene, we used the distribution of branch lengths between genomes known to belong to the same species, and measured each distribution's mean and 95th percentile. The upper threshold was then set by retrieving the highest observed 95th percentile (orange dashed line), and the highest observed mean (magenta dashed line). While the percentage of genes exceeding each threshold varies for each genome, they are relatively consistent, and lead to the same OTU assignment and the same conclusions when investigating tetraploid species. ilAceEphe1 still stands out as possessing more genes which exceed the same-species threshold (no matter what threshold was used) than other genomes.

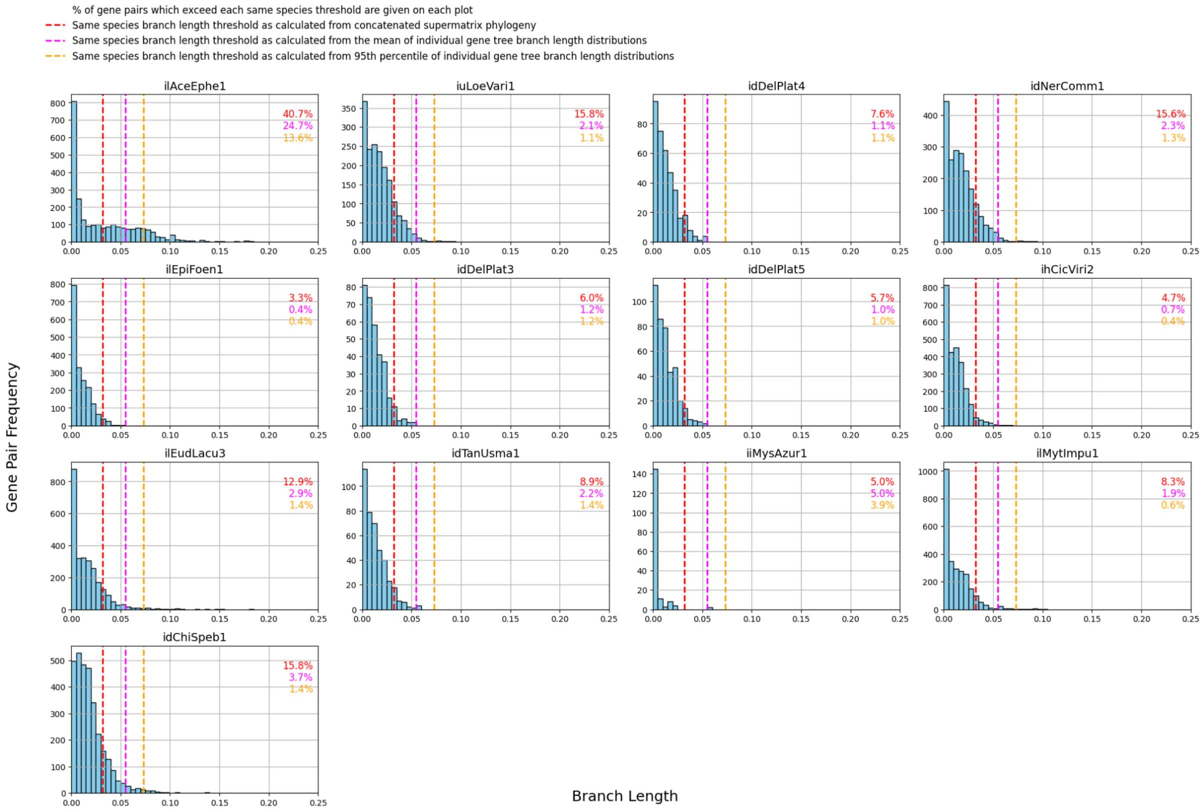


Fig. S2: Relationship between whole-genome phylogeny species delineation thresholds and individual gene phylogeny branch length distribution species delineation thresholds.

We compared our two gene-based metrics (highest 95th percentile and highest mean of branch length distributions of individual gene trees for genomes known to belong to the same species) to the whole-genome-based metric (highest branch length observed between any two same species genomes). We found the relationship between them to be consistent and linear, in line with the fact that they lead to the same conclusions.

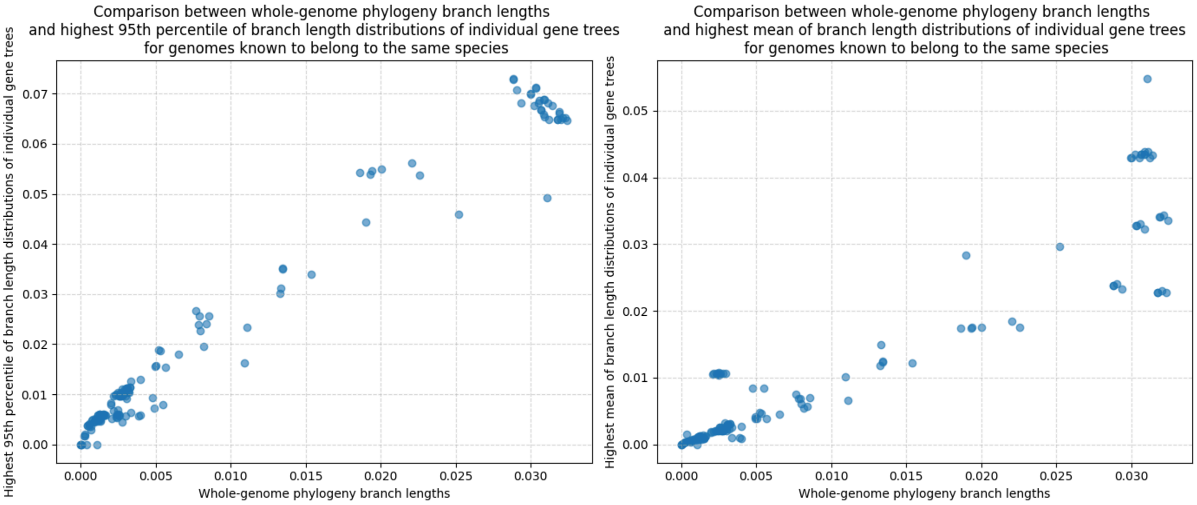
Comparison between whole-genome phylogeny branch lengths
and highest 95th percentile of branch length distributions of individual gene trees
for genomes known to belong to the same species
Highest 95th percentile of branch length distributions of individual gene trees
Whole-genome phylogeny branch lengths
0.000
0.005
0.010
0.015
0.020
0.025
0.030
0.00
0.01
0.02
0.03
0.04
0.05
0.06
0.07
Comparison between whole-genome phylogeny branch lengths
and highest mean of branch length distributions of individual gene trees
for genomes known to belong to the same species
Highest mean of branch length distributions of individual gene trees
Whole-genome phylogeny branch lengths
0.000
0.005
0.010
0.015
0.020
0.025
0.030
0.00
0.01
0.02
0.03
0.04
0.05

### Supporting Information Section 4

Fig. S3: Tetraploid ilAceEphe1.μ is uneven and rearranged.

The number of BUSCO genes found in X haplotypes, along with their total copy number. idChiSpeb1.μ is an even tetraploid, so nearly all its BUSCO genes are in 4 copies, distributed across 4 haplotypes. On the other hand, ilAceEphe1.μ is an uneven tetraploid. The majority of its BUSCO genes are in less than 4 copies, and they are not evenly distributed across its haplotypes. For instance, some BUSCO genes occur in 3 copies present only in a single haplotype.

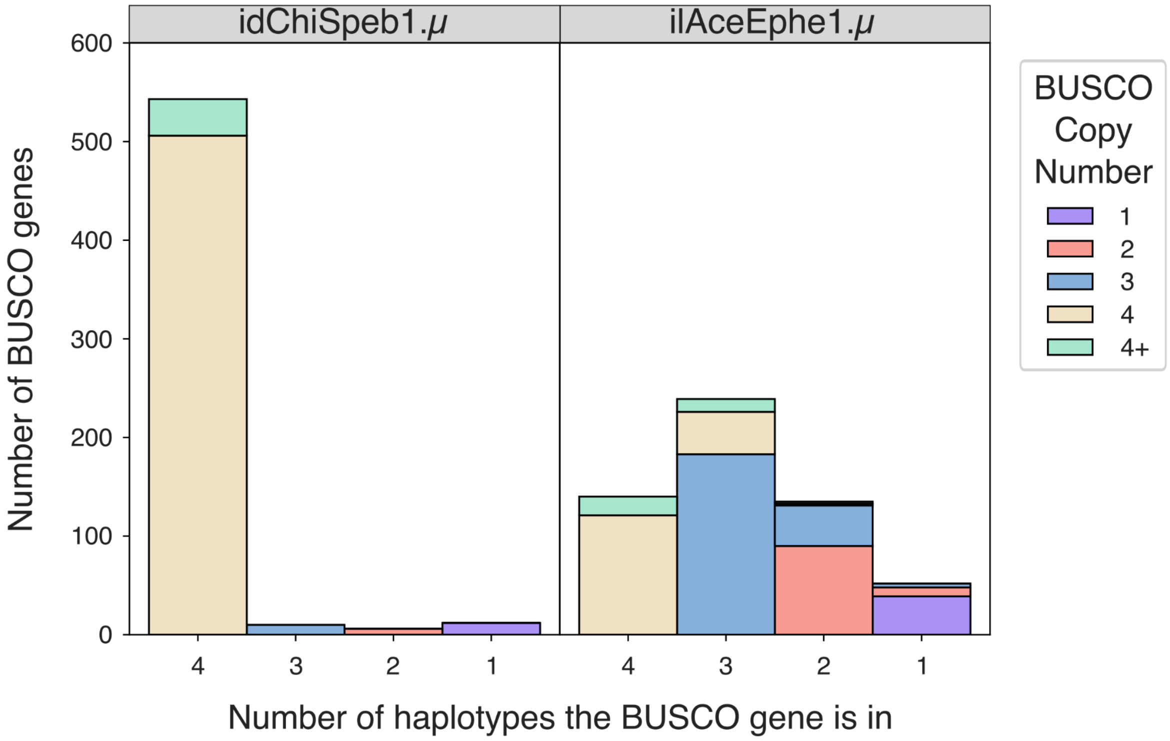

### Supporting Information Section 5

Fig. S4: Phylogeny used by Syngraph, with its internal node labelling.

Each node is labelled with its Syngraph name in a grey box. Yellow boxes indicate the number of chromosomes each genome possesses, and blue boxes indicate the number of chromosomes which possess BUSCO gene markers.

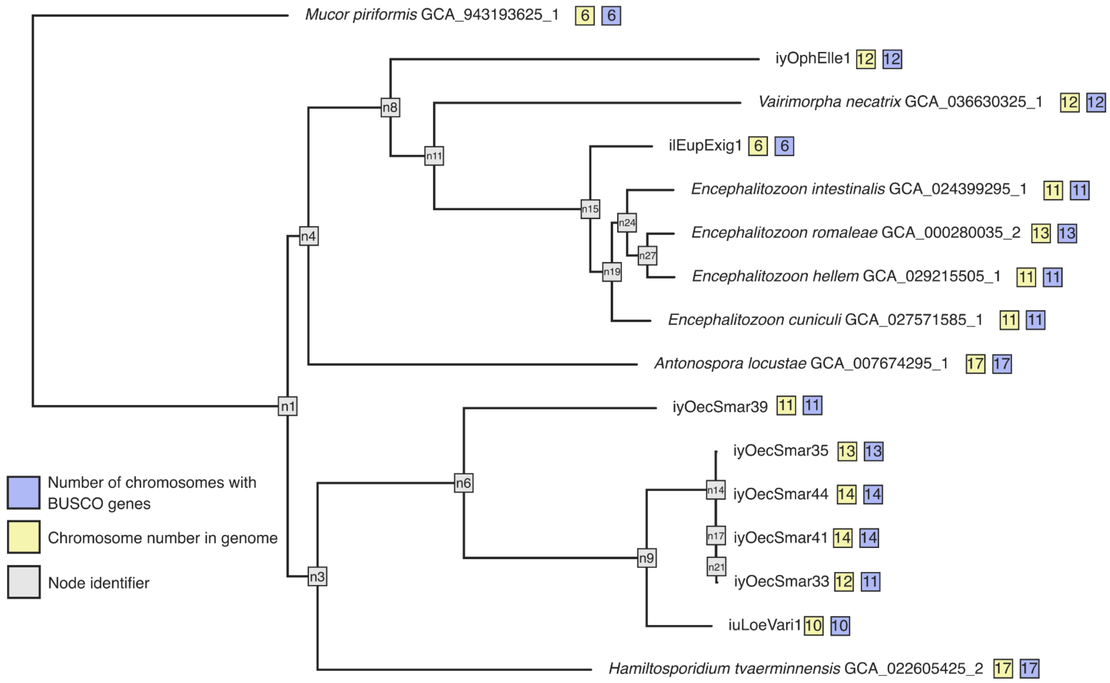


Fig. S5: Number of chromosomes inferred at each node is highly variable.

The number of chromosomes inferred for each node, and the total number of BUSCO genes assigned to a chromosome for each "m". "m" is the parameter in Syngraph to determine the minimum number of genes needed to travel together for the event to be counted as a rearrangement. For example, if m = 3, only rearrangements involving 3 or more genes will be counted. Deep nodes are highly variable and their karyotype (and thus the number of rearrangements that have occurred along each branch) cannot be estimated reliably. See Fig. S4 for node labels on the phylogeny.

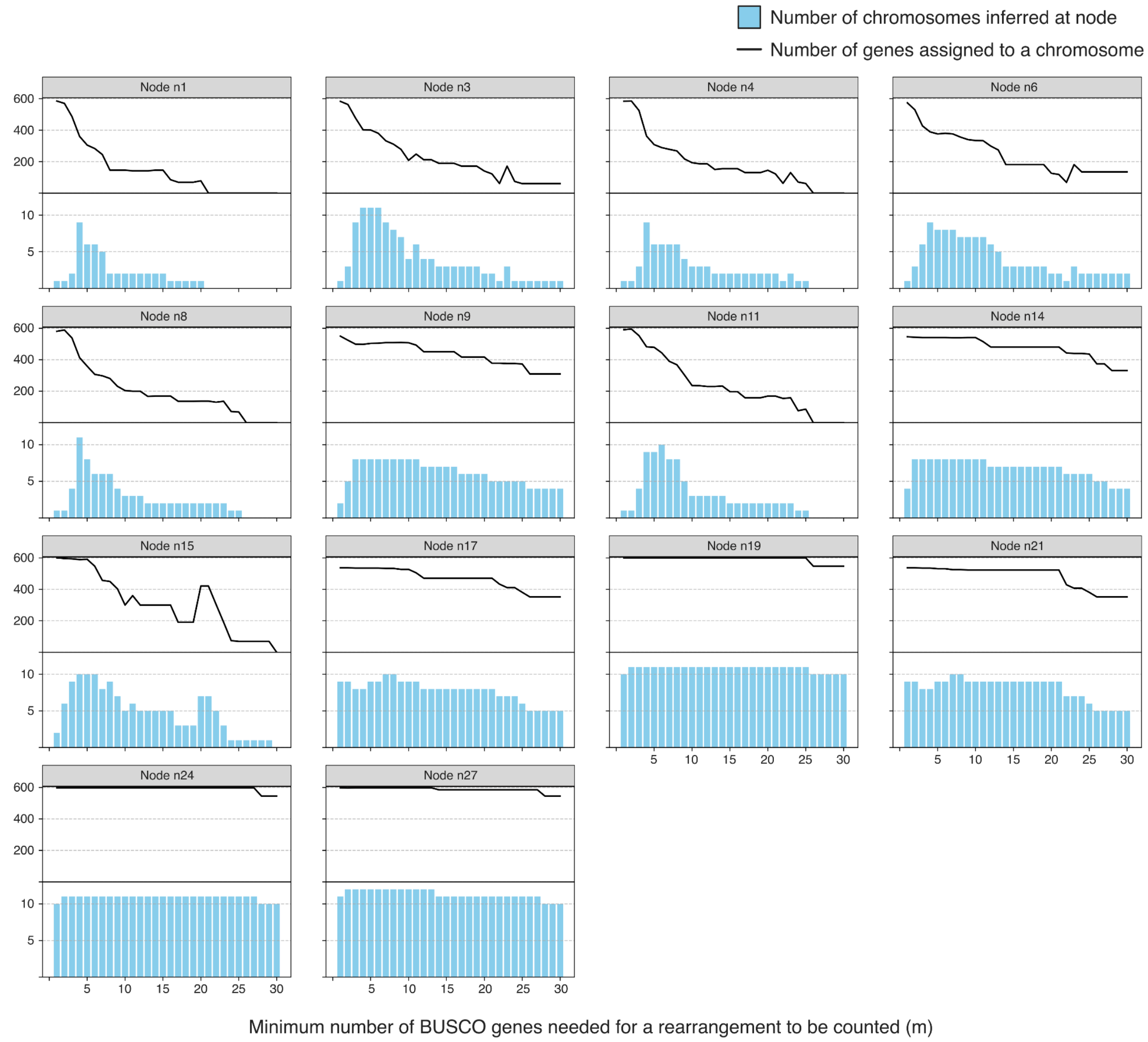


Fig. S6: T-SNE plot depicting BUSCO linkage groups across the microsporidian phylogeny.

Each point represents a BUSCO gene, positioned based on its co-occurrence profile across the chromosome-level microsporidian genomes. Distances between points reflect similarities in co-occurrence. Points are coloured by their assigned chromosome in *Anotonspora locustae*. This disorganised pattern illustrates that the rate of rearrangement is too high for a reliable complete reconstruction of putative ancestral linkage groups. The large-scale patterns are influenced by more densely sampled taxa, see Fig. S7.

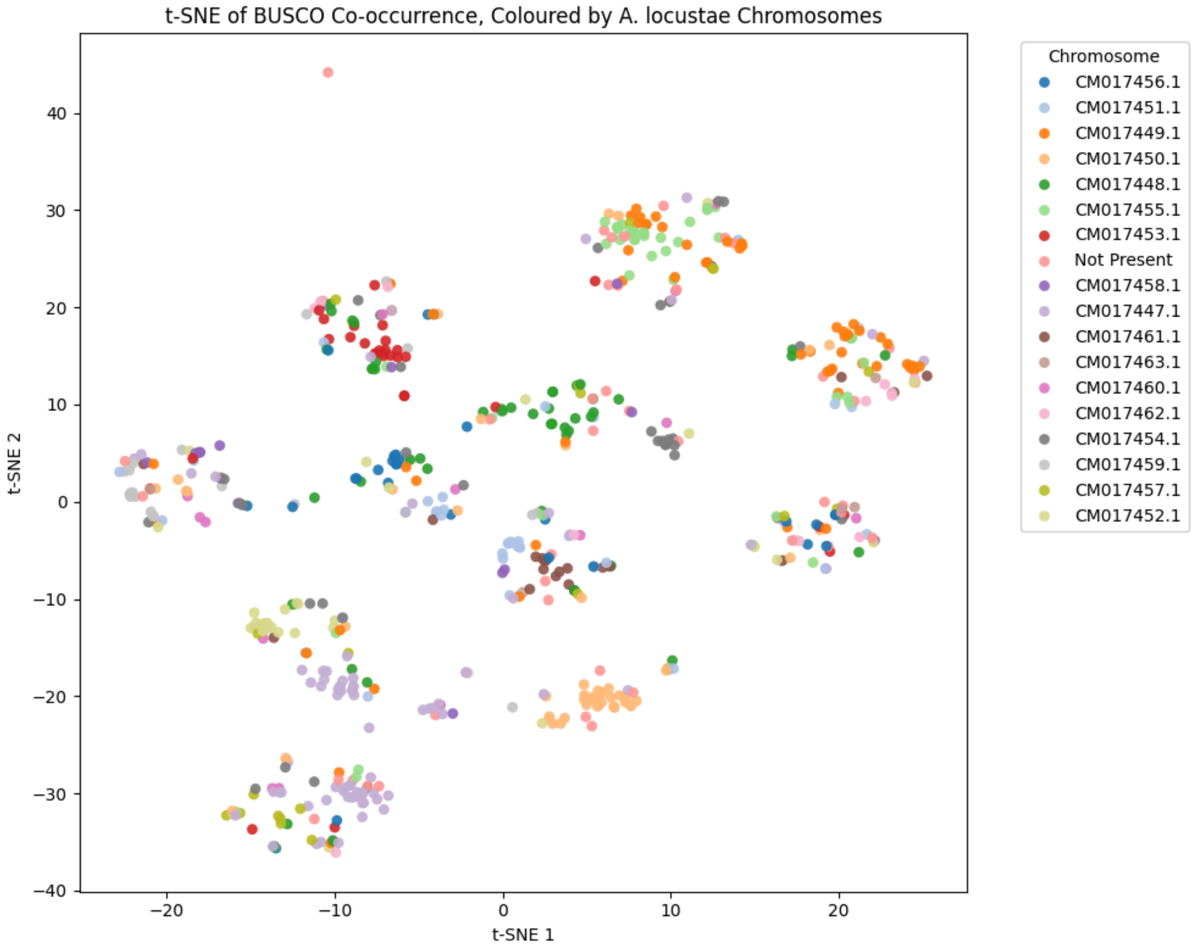


Fig. S7: T-SNE plot depicting BUSCO linkage groups across the microsporidian phylogeny, highlighting clustering influence by more densely sampled taxa.

Each point represents a BUSCO gene, positioned based on its co-occurrence profile across the chromosome-level microsporidian genomes. Distances between points reflect similarities in co-occurrence. Points are coloured by their assigned chromosome in *Encephalitozoon cuniculi*. This disorganised pattern illustrates that the rate of rearrangement is too high for a reliable complete reconstruction of putative ancestral linkage groups. The large-scale patterns are influenced by more densely sampled taxa, such as *Encephalitozoon cuniculi*.

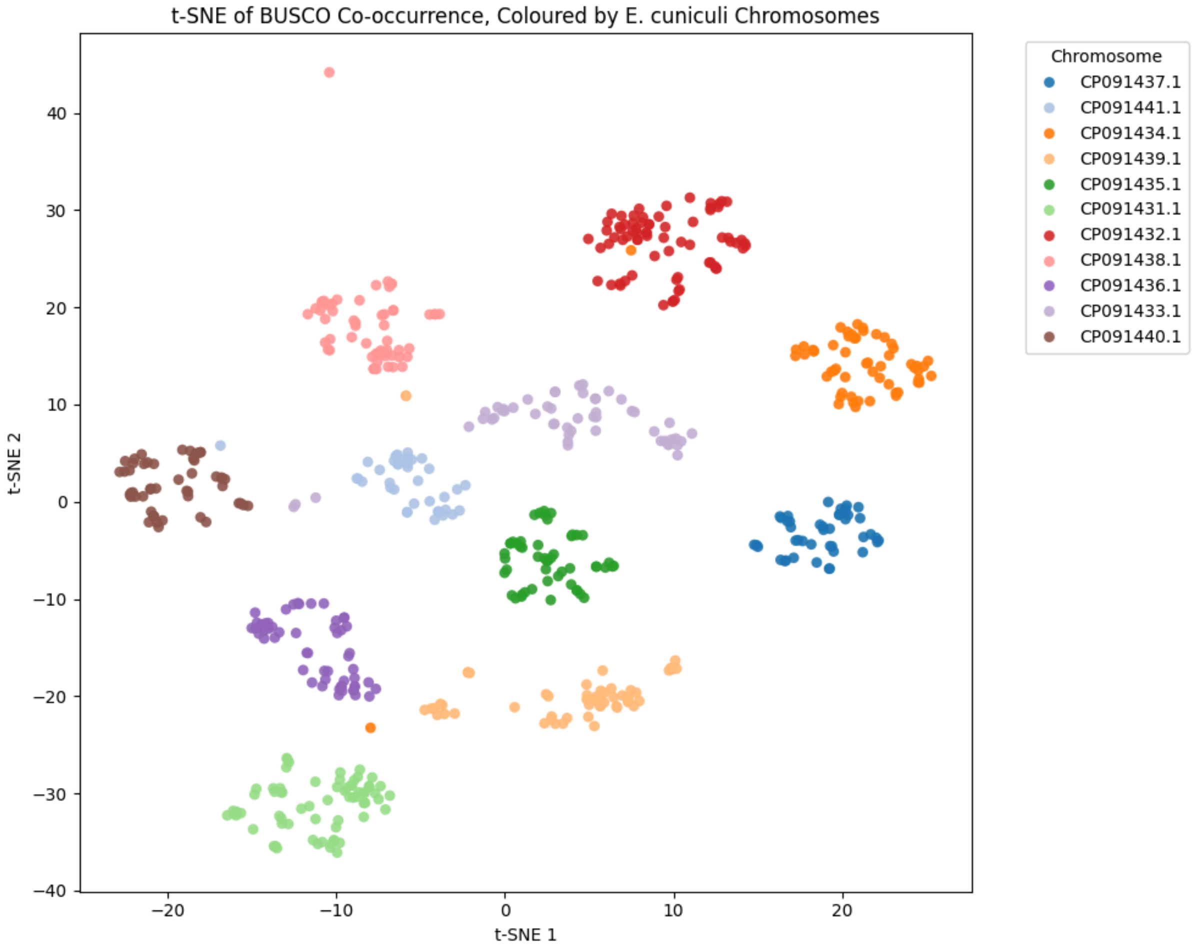


Fig. S8: Synteny plots of chromosomal microsporidian genome assemblies.

Genome-wide synteny plots of all available chromosomal microsporidian genome assemblies. Each line represents a single-copy BUSCO (microsporidia_odb10) [9]. BUSCOs are painted by their chromosomal position in *A. locustae*. Plot was generated by modifying ribbon plot scripts from https://github.com/conchoecia/odp [10].

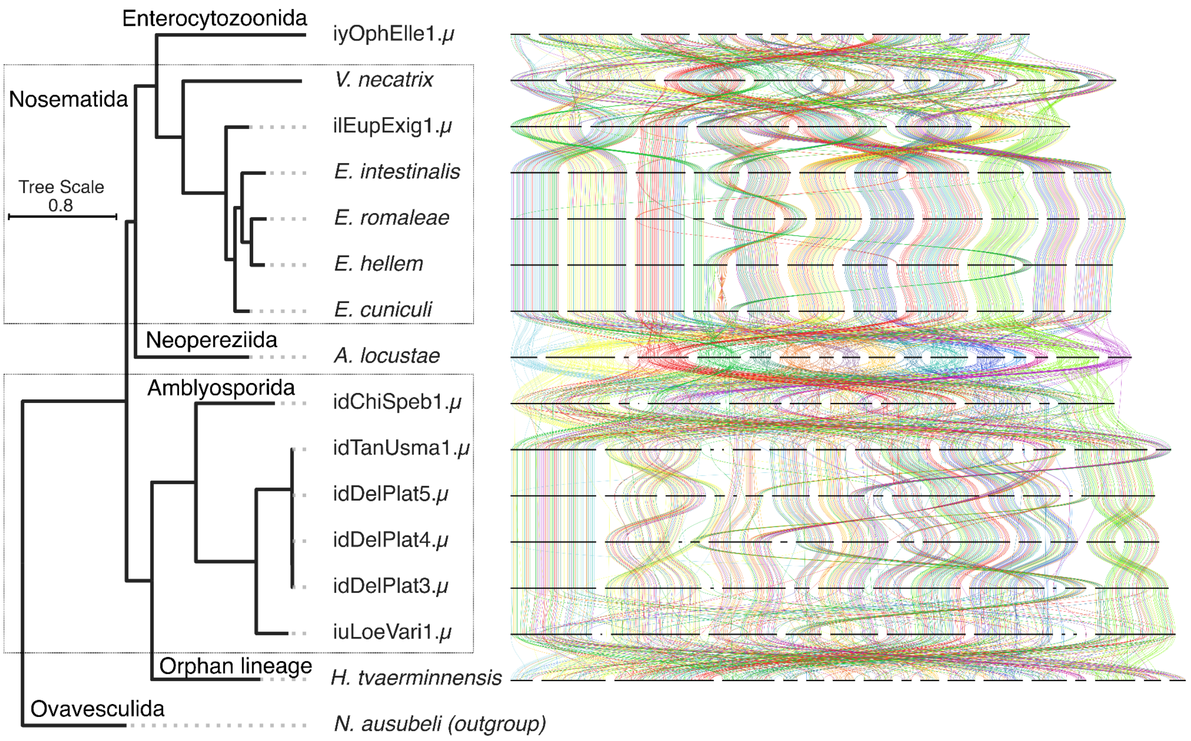


Fig. S9: Synteny plots of chromosomal microsporidian genome assemblies.

Genome-wide synteny plots of all available chromosomal microsporidian genome assemblies. Each line represents a single-copy BUSCO (microsporidia_odb10) [9]. BUSCOs are painted by their chromosomal position in *H. tvaerminnensis*. Plot was generated by modifying ribbon plot scripts from https://github.com/conchoecia/odp [10].

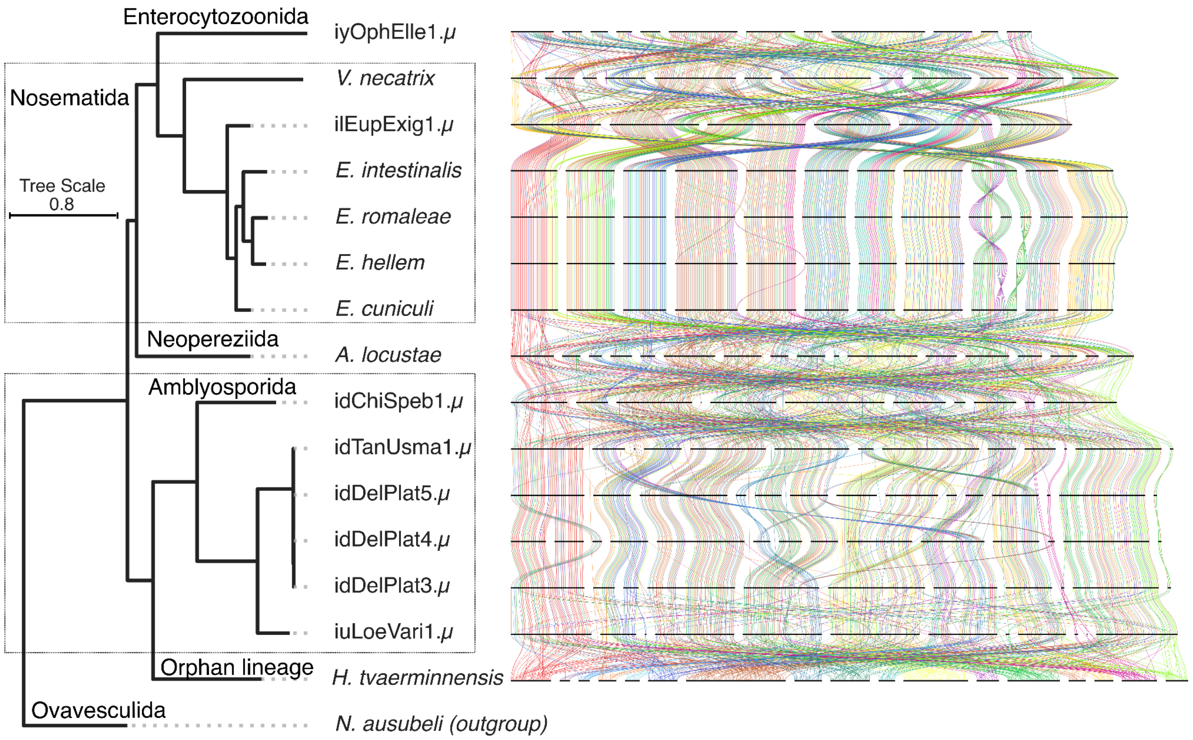

Enterocytozoonida
iyOphElle1.μ
Nosematida
V. necatrix
ilEupExig1.μ
Tree Scale
0.8
E. intestinalis
E. romaleae
E. hellem
E. cuniculi
Neopereziida
A. locustae
Amblyosporida
idChiSpeb1.μ
idTanUsma1.μ
idDelPlat5.μ
idDelPlat4.μ
idDelPlat3.μ
iuLoeVari1.μ
Orphan lineage
H. tvaerminnensis
Ovavesculida
N. ausubeli (outgroup)

### References


1. Challis R, Richards E, Rajan J, Cochrane G, Blaxter M. BlobToolKit – Interactive Quality Assessment of Genome Assemblies. G3 Genes|Genomes|Genetics. 2020;10: 1361–1374. doi:10.1534/g3.119.400908

2. Ranallo-Benavidez TR, Jaron KS, Schatz MC. GenomeScope 2.0 and Smudgeplot for reference-free profiling of polyploid genomes. Nat Commun. 2020;11: 1–10. doi:10.1038/s41467-020-14998-3

3. Marçais G, Kingsford C. A fast, lock-free approach for efficient parallel counting of occurrences of k-mers. Bioinformatics. 2011;27: 764–770. doi:10.1093/bioinformatics/btr011

4. PretextView: OpenGL Powered Pretext Contact Map Viewer. Github; Available: https://github.com/sanger-tol/PretextView

5. Flynn JM, Hubley R, Goubert C, Rosen J, Clark AG, Feschotte C, et al. RepeatModeler2 for automated genomic discovery of transposable element families. Proceedings of the National Academy of Sciences. 2020;117: 9451–9457. doi:10.1073/pnas.1921046117

6. Seemann T. Prokka: rapid prokaryotic genome annotation. Bioinformatics. 2014;30: 2068–2069. doi:10.1093/bioinformatics/btu153

7. Smit AFA, Hubley R, Green P. RepeatMasker Open-4.0.2013-2015. 2013. Available: <http://www.repeatmasker.org>

8. Pagel M. Detecting correlated evolution on phylogenies: a general method for the comparative analysis of discrete characters. Proceedings of the Royal Society of London Series B: Biological Sciences. 1994;255: 37–45. doi:10.1098/rspb.1994.0006

9. Simão FA, Waterhouse RM, Ioannidis P, Kriventseva EV, Zdobnov EM. BUSCO: assessing genome assembly and annotation completeness with single-copy orthologs. Bioinformatics. 2015;31: 3210–3212. doi:10.1093/bioinformatics/btv351

10. Schultz DT, Haddock SHD, Bredeson JV, Green RE, Simakov O, Rokhsar DS. Ancient gene linkages support ctenophores as sister to other animals. Nature. 2023;618: 110–117. doi:10.1038/s41586-023-05936-6
